## Supplementary Information for "CiFi: Accurate long-read chromosome conformation capture with low-input requirements"

**1. Supplementary Figures**

**Supplementary Figure 1.** Genome-wide sequencing coverage statistics.
**Supplementary Figure 2.** Workflow of the CiFi library preparation protocol.
**Supplementary Figure 3.** Proportion of segments within a CiFi read mapping to the same chromosome.
**Supplementary Figure 4.** Mapping quality comparison of CiFi, Hi-C, and Pore-C reads across the genome.
**Supplementary Figure 5.** Normalized read coverage of CiFi, Pore-C, and Hi-C across the genome.
**Supplementary Figure 6.** Comparisons of chromatin contacts for human LCL GM12878 between DpnII CiFi and Hi-C.
**Supplementary Figure 7.** Correlation of chromatin contacts between DpnII CiFi and Hi-C.
**Supplementary Figure 8.** Topologically associating domains for GM2878 LCL DpnII CiFi vs Hi-C across a unique space.
**Supplementary Figure 9.** Comparison of chromatin contacts for human LCL GM12878 between DpnII CiFi and Pore-C genome-wide and across chromosomes 1-5.
**Supplementary Figure 10.** Correlation of chromatin contact signal at 2.5 Mbp resolutions between DpnII CiFi and Pore-C.
**Supplementary Figure 11**. Comparisons of chromatin contacts for human LCL GM12878 across different cell titrations for chromosome 1.
**Supplementary Figure 12.** Percentage of reads mapped and normalized read coverage of CiFi and Hi-C across the genome for *Anopheles coluzzii*.
**Supplementary Figure 13.** Scaffolding the Mediterranean fruit fly assembly with CiFi.
**Supplementary Figure 14**: Scaffolding the Mediterranean fruit fly assembly with varying coverages of CiFi segments.
**Supplementary Figure 15:** Scaffolding the Mediterranean fruit fly assembly with varying coverages of Hi-C reads.
**Supplementary Figure 16:** Comparison of CiFi and Hi-C datasets across sex chromosomes mapped to the new Mediterranean fruit fly genome assembly.
**Supplementary Figure 17.** Quality check of 3C DNA.
**Supplementary Figure 18.** Verification of fragment size during CiFi library preparation.

**2. Supplementary Tables**

**Supplementary Table 1.** Comparison of Pore-C and Multi-Contact 3C protocols.
**Supplementary Table 2.** Contact signal correlation for GM12878.
**Supplementary Table 3.** Genome-wide TopDom TAD measure of concordance.
**Supplementary Table 4.** Genome-wide TopDom TAD Jaccard Index.
**Supplementary Table 5.** Correlation of CiFi chromatin contacts across cell titration.
**Supplementary Table 6.** Coordinates of AcolN3 repeat annotations.
**Supplementary Table 7.** *An. coluzzii* mosquito chromosome coverage at MAPQ1.
**Supplementary Table 8.** Contact signal correlation for *An. coluzzii* mosquito between CiFi and Hi-C.
**Supplementary Table 9.** Mediterranean fruit fly assembly statistics.

**3. Extended Experimental procedures: Detailed CiFi Protocol**

**Part 1: 3C library preparation**

i. Cross-linking and quenching.
ii. Restriction enzyme digestion.
iii. Proximity ligation and reverse cross-linking.
iv. Protein degradation and DNA purification.

**Part 2: SMRTbell library preparation from modified ultra-low DNA input**

**Part 2A: SMRTbell library preparation with Express Template Prep Kit 2.0**

i. Removing single-strand overhangs.
ii. Repair DNA damage.
iii. Repair ends/A-tailing.
iv. Adapter ligation.
v. Purification of SMRTbell library.
vi. Library amplification by modified PCR.
vii. Purification of amplified DNA.
viii. Repair DNA damage.
ix. Repair ends/A-tailing.
x. Adapter ligation.
xi. Purification of SMRTbell library.
xii. BluePippin or diluted AMPure PB bead cleanup, size selection, and sequencing.

**Part 2B: SMRTbell library preparation with Prep Kit 3.0**

i. Repair and A-tailing of digested 3C DNA.
ii. Ligation of linear amplification adapter and cleanup.
iii. Amplification and cleanup.
iv. Repair and A-tailing of amplified DNA.
v. SMRTbell adapter ligation and cleanup.
vi. Nuclease treatment.
vii. BluePippin or diluted AMPure PB bead cleanup, size selection, and sequencing.

**4. Uncropped Gel Image:** Supplementary Figure 17

**5. Supplementary References**

### 1. Supplementary Figures

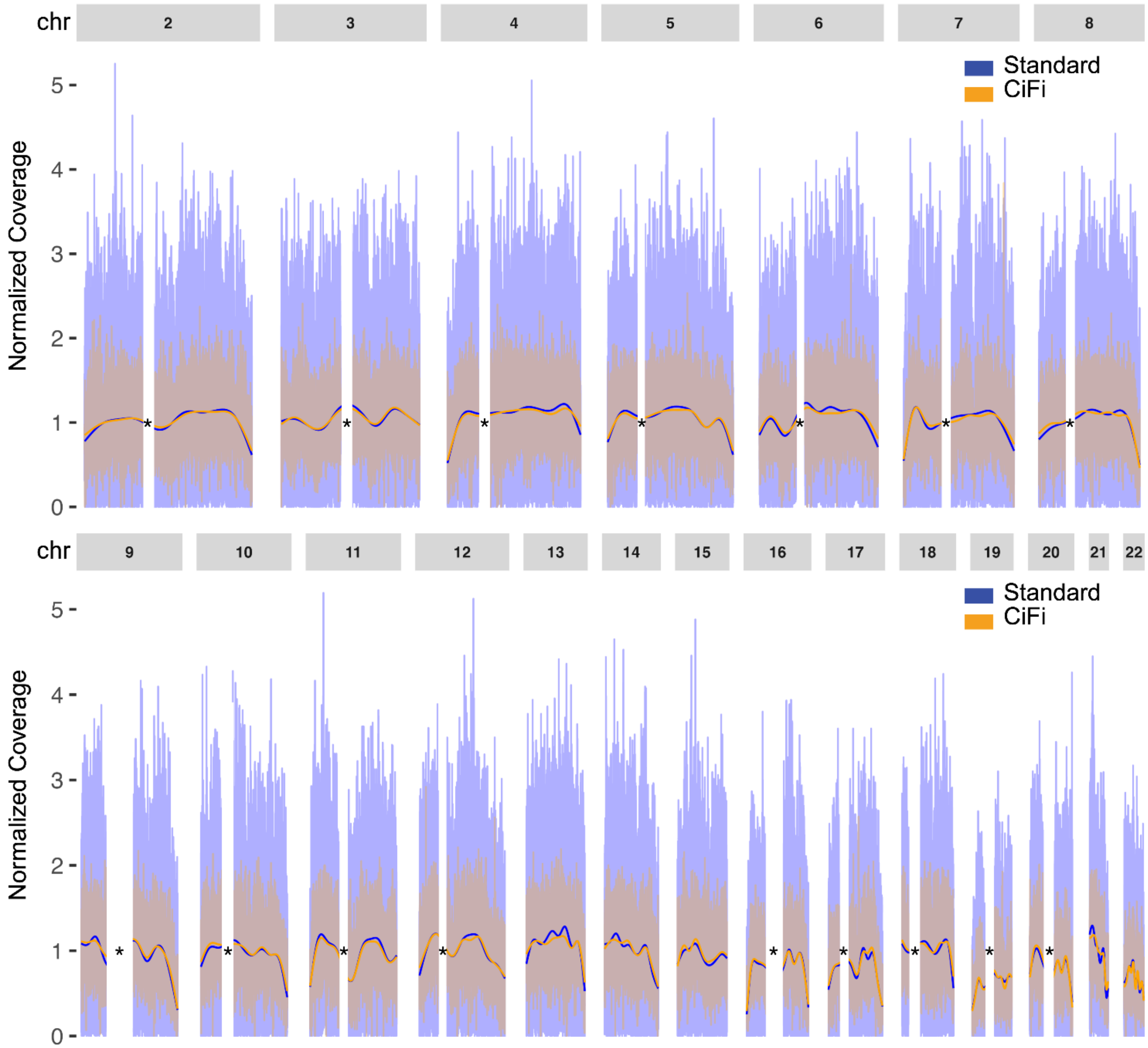

**Supplementary Figure 1. Genome-wide sequencing coverage statistics.** Normalized read-coverage comparison of Sequel II data for DpnII 3C libraries generated without (Pore-C/Standard) and with the amplification-protocol (CiFi) for GM12878 across human chromosomes 2 through 22 *excluding non-unique regions. Source data are provided as a Source Data file.

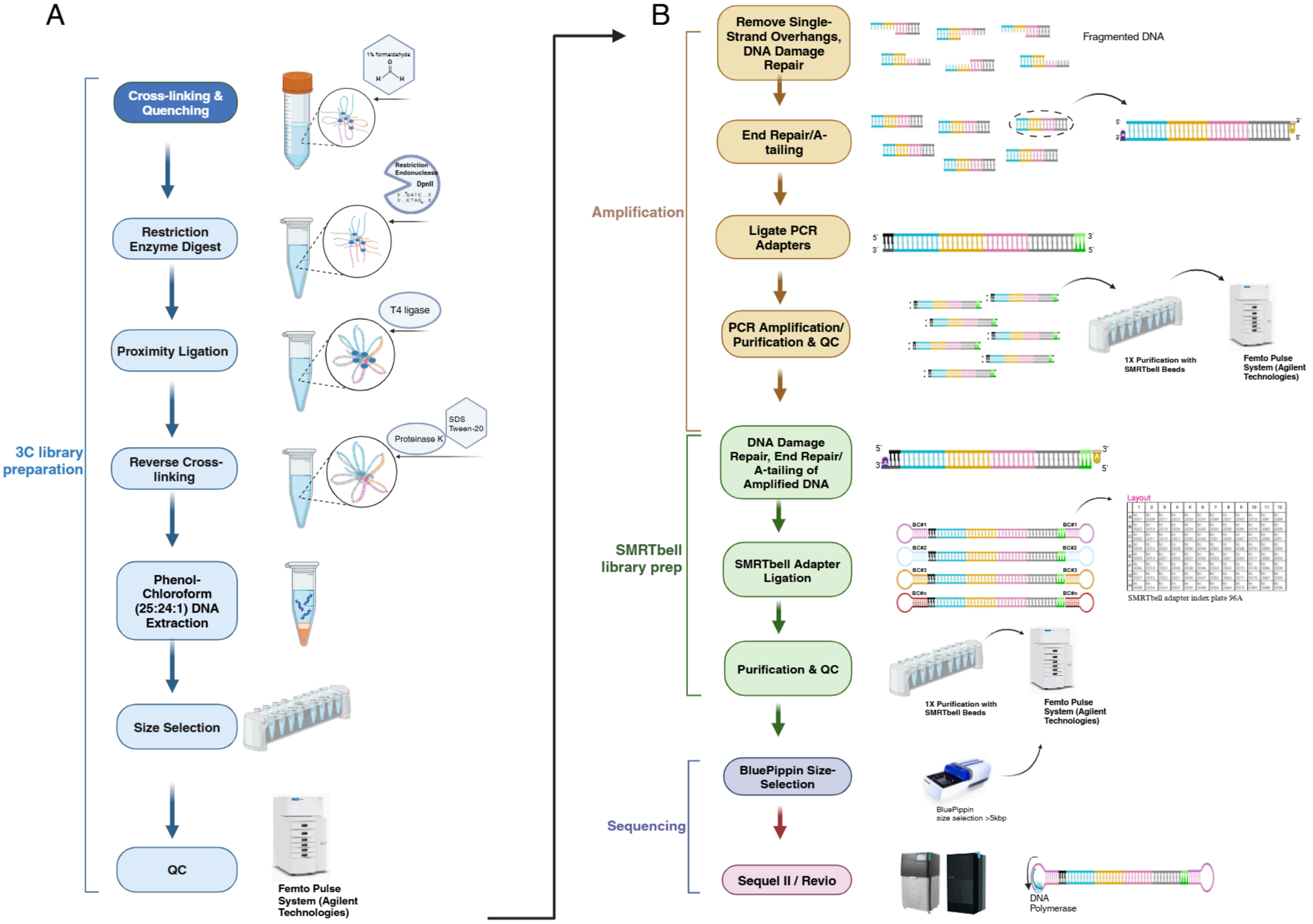

**Supplementary Figure 2. Workflow of the CiFi library preparation protocol.** For generating HiFi sequencing libraries from 3C DNA, the workflow is divided into two main parts: **(A)** 3C Library Preparation: Includes cross-linking chromatin, restriction enzyme digestion, proximity ligation, reverse cross-linking, DNA purification, and size selection. **(B)** SMRTbell Library Preparation: Following amplification, DNA damage repair, A-tailing, and adapter ligation are performed. Final libraries undergo size selection using BluePippin and sequencing on the PacBio Sequel II or Revio systems. Created in BioRender. Dennis, M. (2025) https://BioRender.com/97iyp0n.

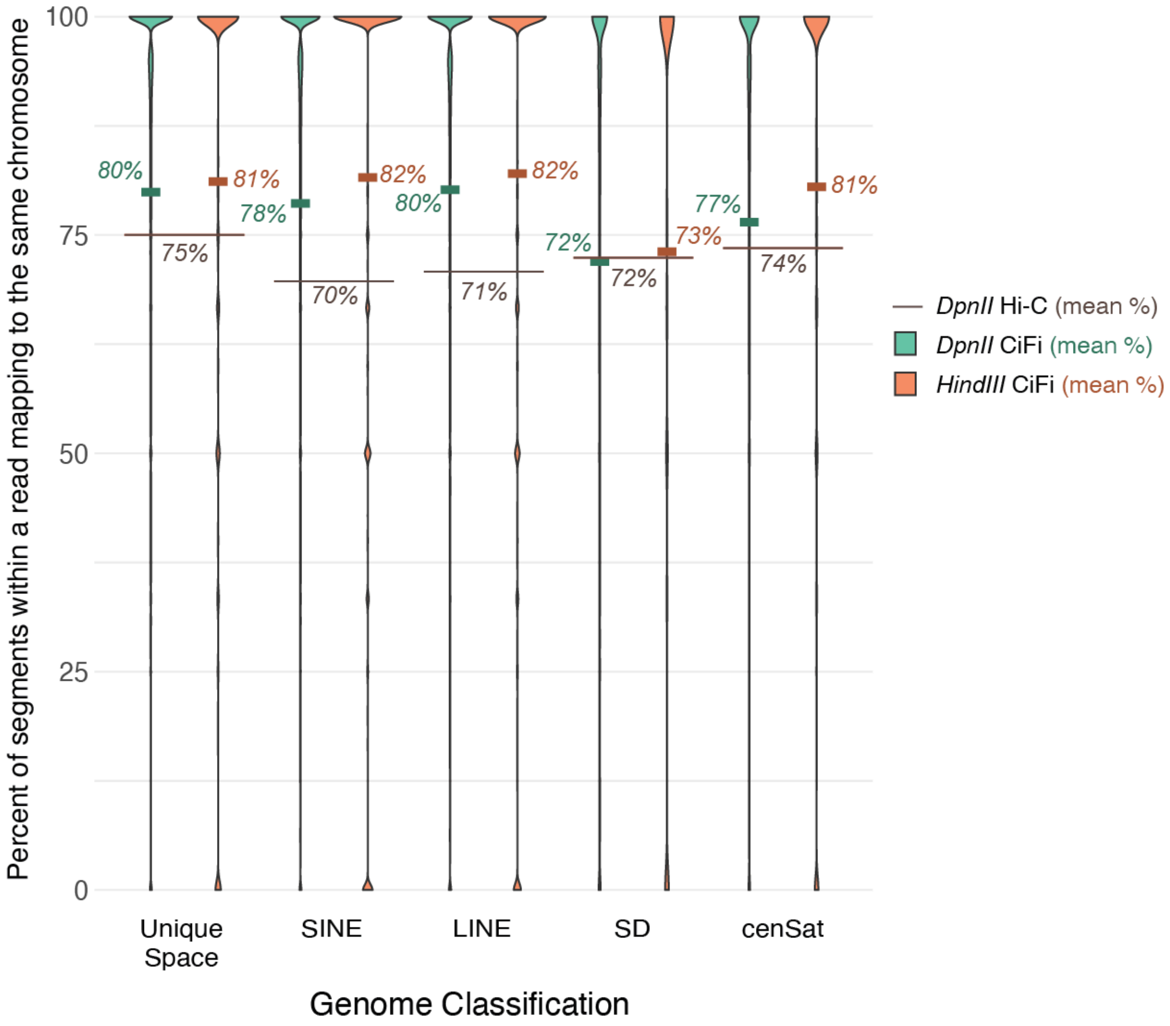

**Supplementary Figure 3. Proportion of segments within a CiFi read mapping to the same chromosome.** The percentage of segments within a CiFi read mapping to the same chromosome, per segment, depicted as violin plots. The segments are characterized by the genomic bin that they share overlap with. Solid lines represent the mean for each region/restriction enzyme group with the gray line representing DpnII Hi-C average. Only the average was calculated for Illumina reads because analysis of paired ends produces a bimodal result (0% or 100%). Source data are provided through NCBI BioProject accession PRJEB83708.

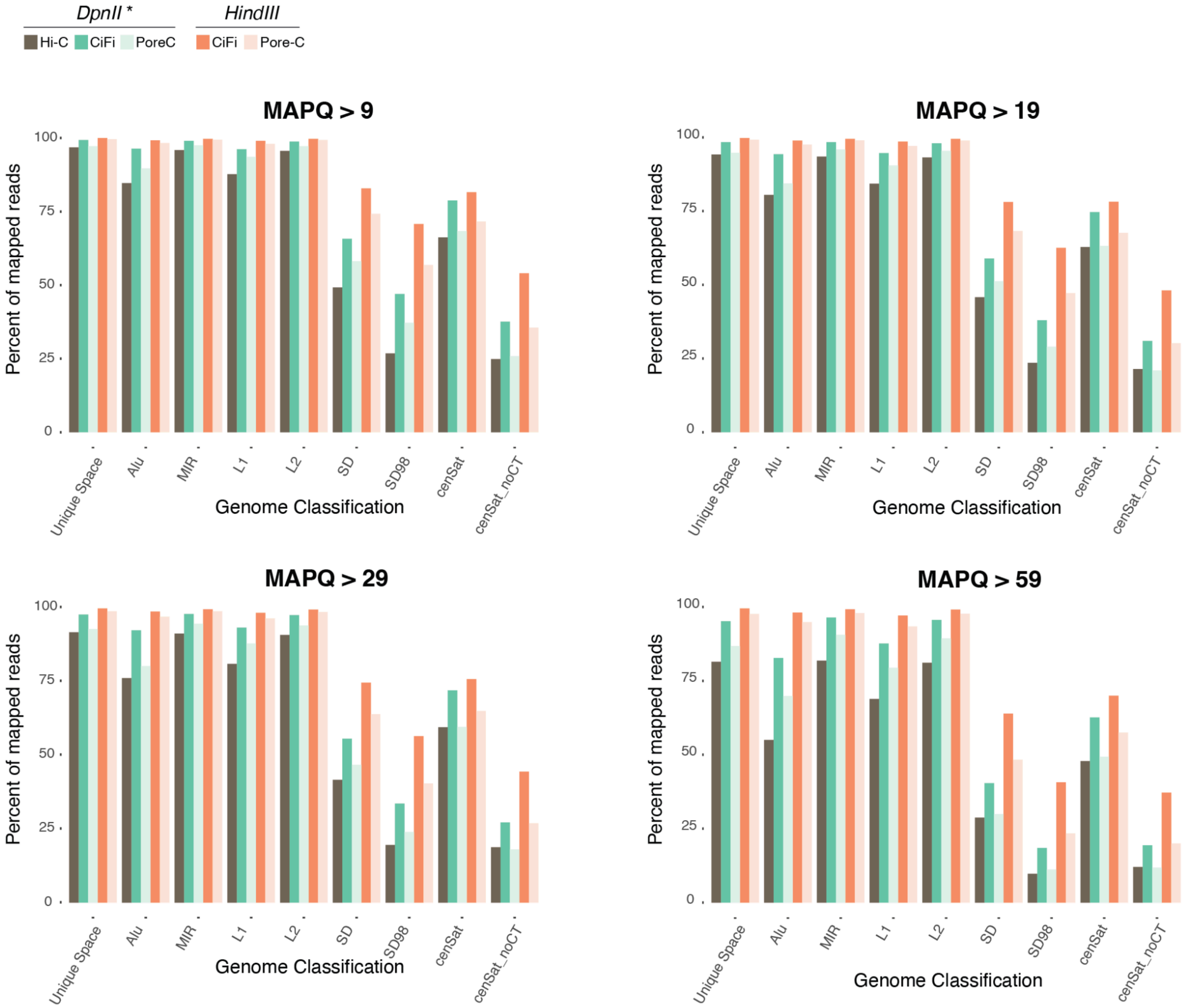

**Supplementary Figure 4. Mapping quality comparison of CiFi, Hi-C, and Pore-C reads across the genome.** Percentage of reads (segments for CiFi and Pore-C) with varied MAPQ cutoffs (indicated above each plot) for Hi-C with Illumina [1], CiFi, and Pore-C [2] across different repetitive genome classifications (short interspersed nuclear elements (SINEs) *Alu* and *Mir*, long interspersed nuclear elements (LINEs) *L1* and *L2*, segmental duplications at both 90% (SD) and 98% (SD98) identity, and centromeres with and without the centromeric transition (CT) regions. Source data are provided as a Source Data file.

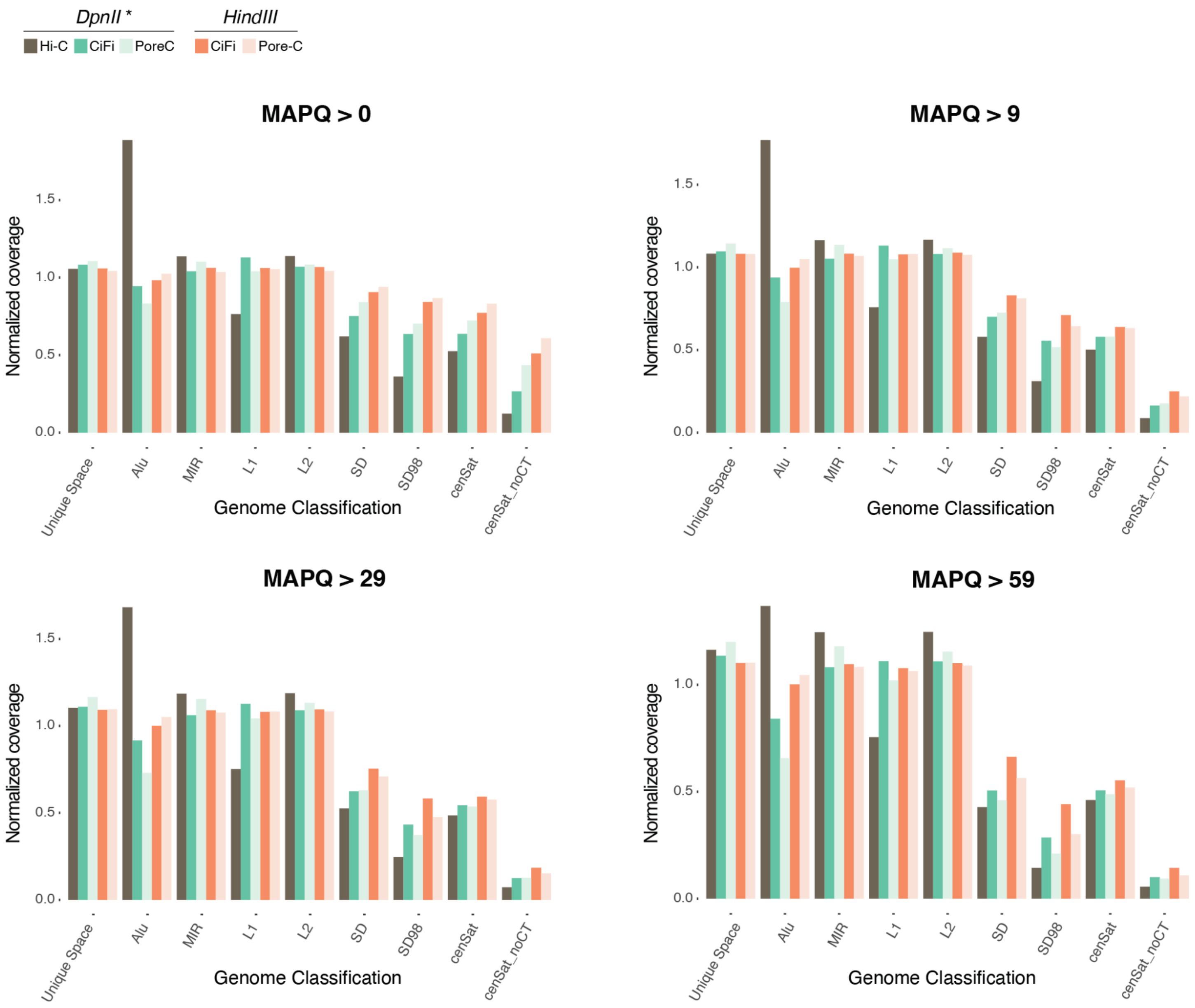

**Supplementary Figure 5. Normalized read coverage of CiFi, Pore-C, and Hi-C across the genome.** Read coverage ratio (ratio of coverage per region to genomewide coverage) with varied MAPQ cutoffs (indicated above each plot) for Hi-C with Illumina [1], CiFi, and Pore-C [2] across different repetitive genome classifications (short interspersed nuclear elements (SINEs) Alu and Mir, long interspersed nuclear elements (LINEs) L1 and L2, segmental duplications at both 90% (SD) and 98% (SD98) identity, and centromeres with and without the centromeric transition (CT) regions) divided by the genome-wide coverage for each library. Source data are provided as a Source Data file.

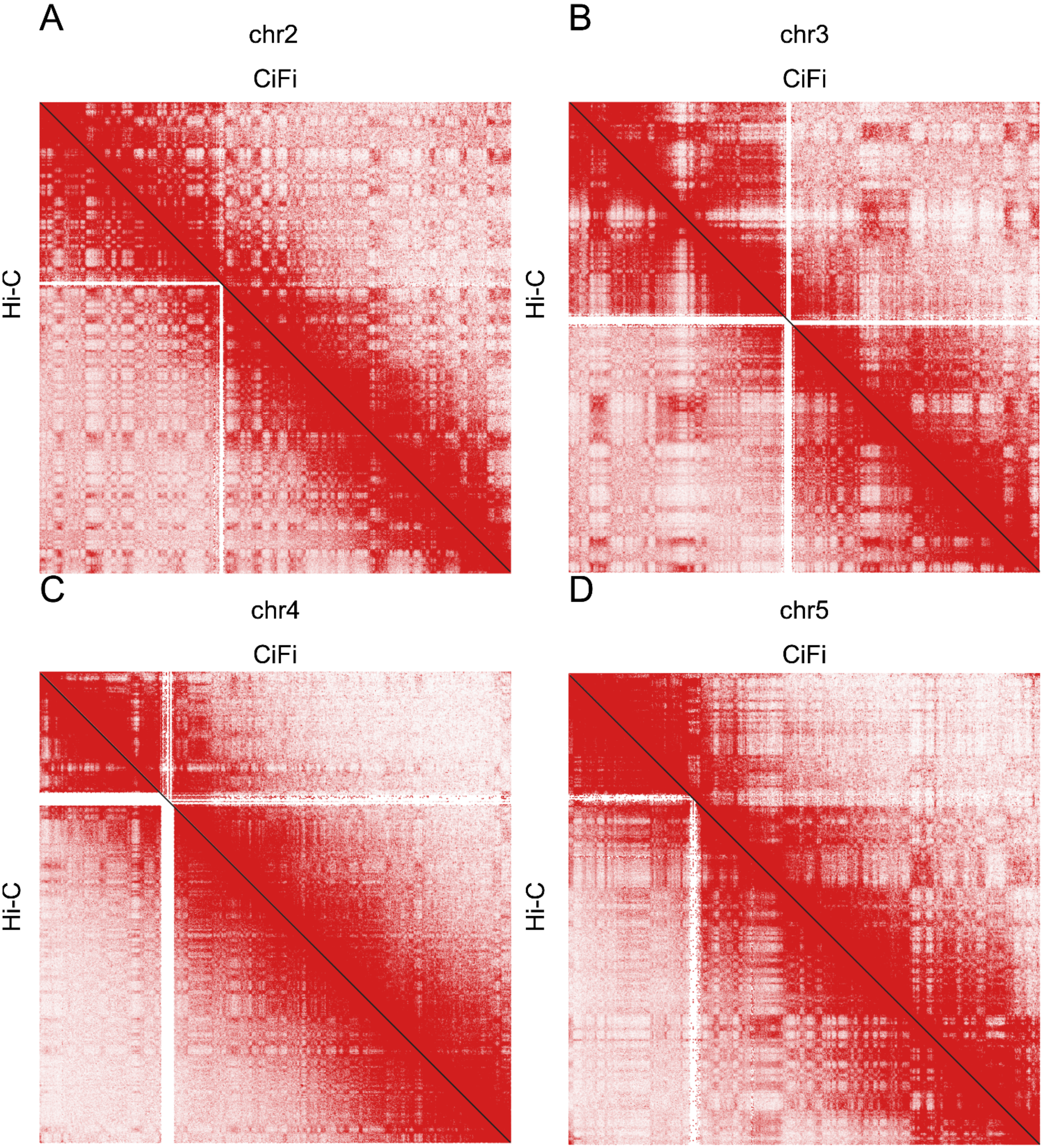

**Supplementary Figure 6. Comparisons of chromatin contacts for human LCL GM12878 between DpnII CiFi and Hi-C.** Chromosome-scale pairwise interaction maps at 250-kbp resolution for **(A)** chromosome 2, **(B)** chromosome 3, **(C)** chromosome 4, and **(D)** chromosome 5. Red color scales with the number of paired reads per bin. Contact matrices are normalized using Knight–Ruiz algorithm. CiFi is above the diagonal while Hi-C is below. Source data are provided through NCBI BioProject accession PRJEB83708.

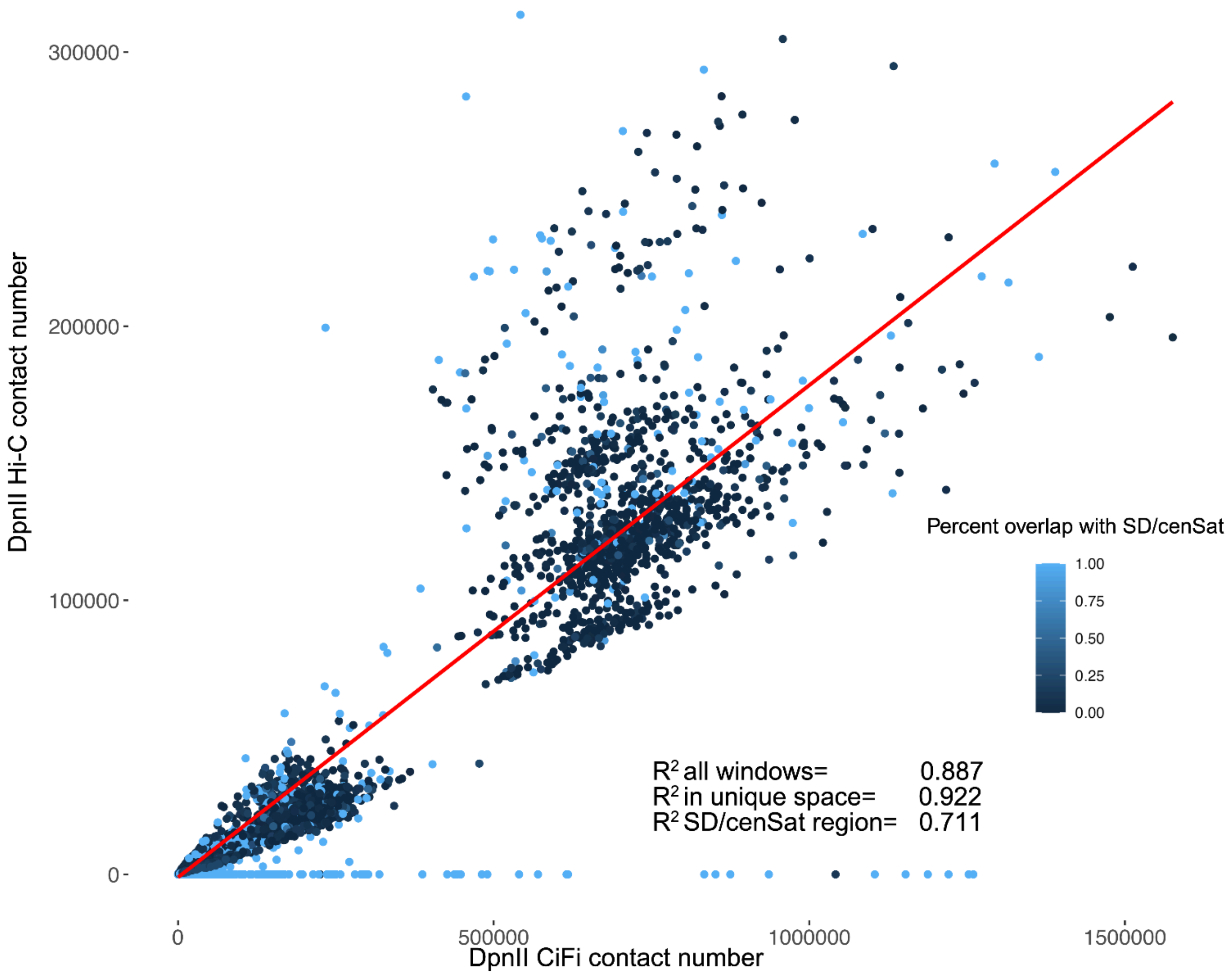

**Supplementary Figure 7. Correlation of chromatin contacts between DpnII CiFi and Hi-C.** Analyzing the $R^2$ across all windows at 2.5-Mbp resolution, those in unique space (< 50% overlap with SD/cenSat), or those in SD/cenSat space (>50% overlap), we observe the highest correlation when only accounting for windows in unique space and the lowest when looking at windows that overlap SD/cenSat. This discordance shows that CiFi leads to a gain of contacts across repetitive regions. Source data are provided as a Source Data file.

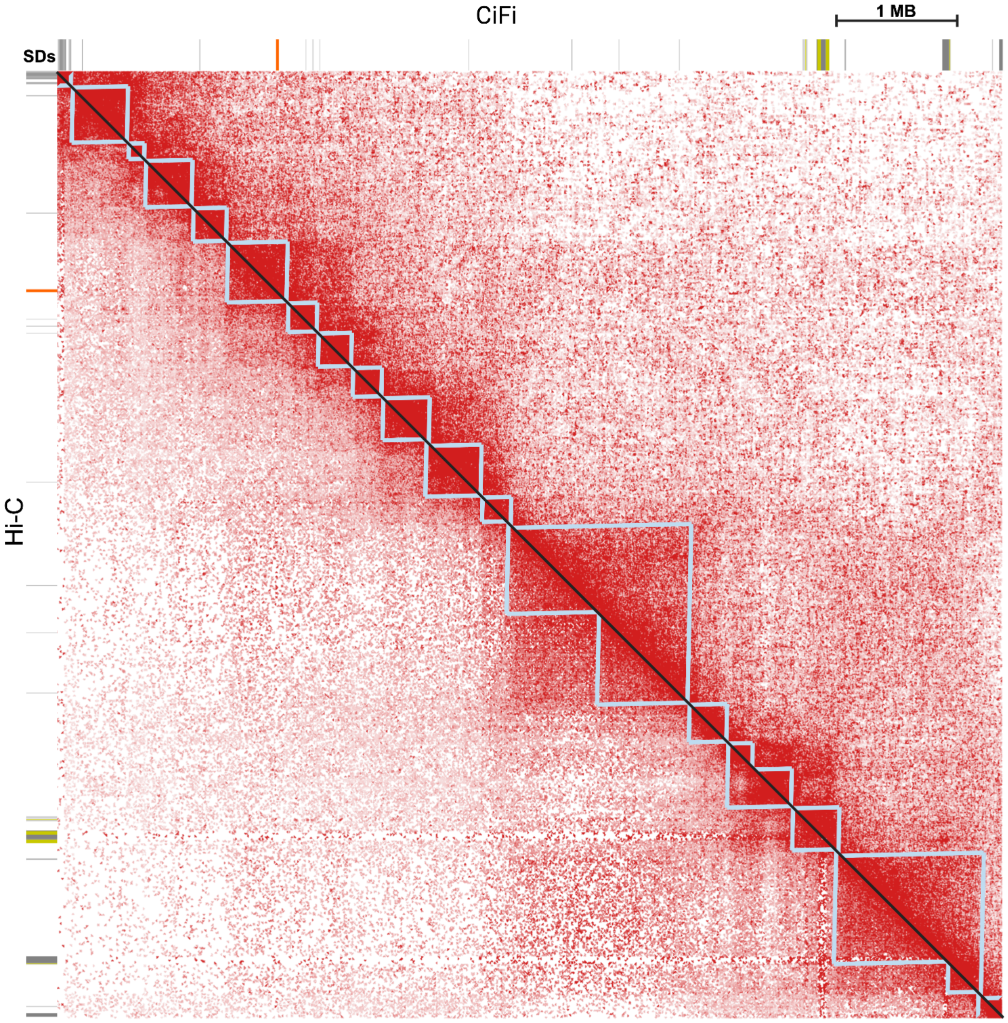

**Supplementary Figure 8. Topologically associating domains for GM2878 LCL DpnII CiFi vs Hi-C across a unique space.** Human chromosome 2 (chr2:98,000,000-109,000,000, T2T-CHM1_v2; 50-kbp resolution), contacts are normalized using Knight–Ruiz algorithm. Topologically associating domains are represented as light blue triangles. CiFi is above the diagonal while Hi-C is below. Source data are provided through NCBI BioProject accession PRJEB83708.

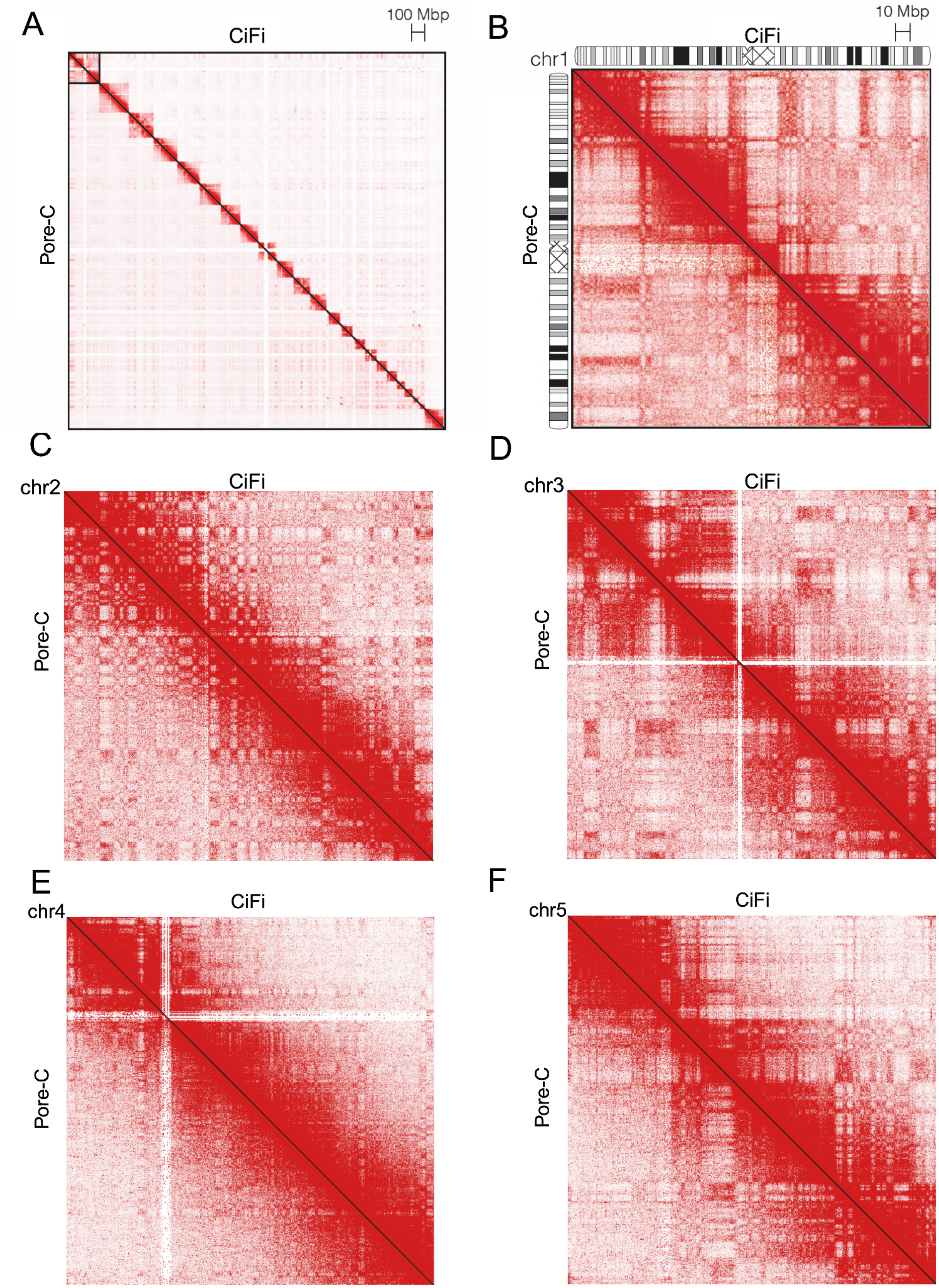

**Supplementary Figure 9. Comparison of chromatin contacts for human LCL GM12878 between DpnII CiFi and Pore-C genome wide and across chromosomes 1-5.** Chromosome-scale pairwise interaction maps at 250-kbp resolution for **(A)** genome wide **(B)** chromosome 1, **(B)** chromosome 2, **(C)** chromosome 3, **(D)** chromosome 4, and **(E)** chromosome 5. Red color scales with the number of paired reads per bin. Contact matrices are normalized using Knight–Ruiz algorithm. CiFi is above the diagonal while Pore-C is below. Source data are provided through NCBI BioProject accession PRJEB83708.

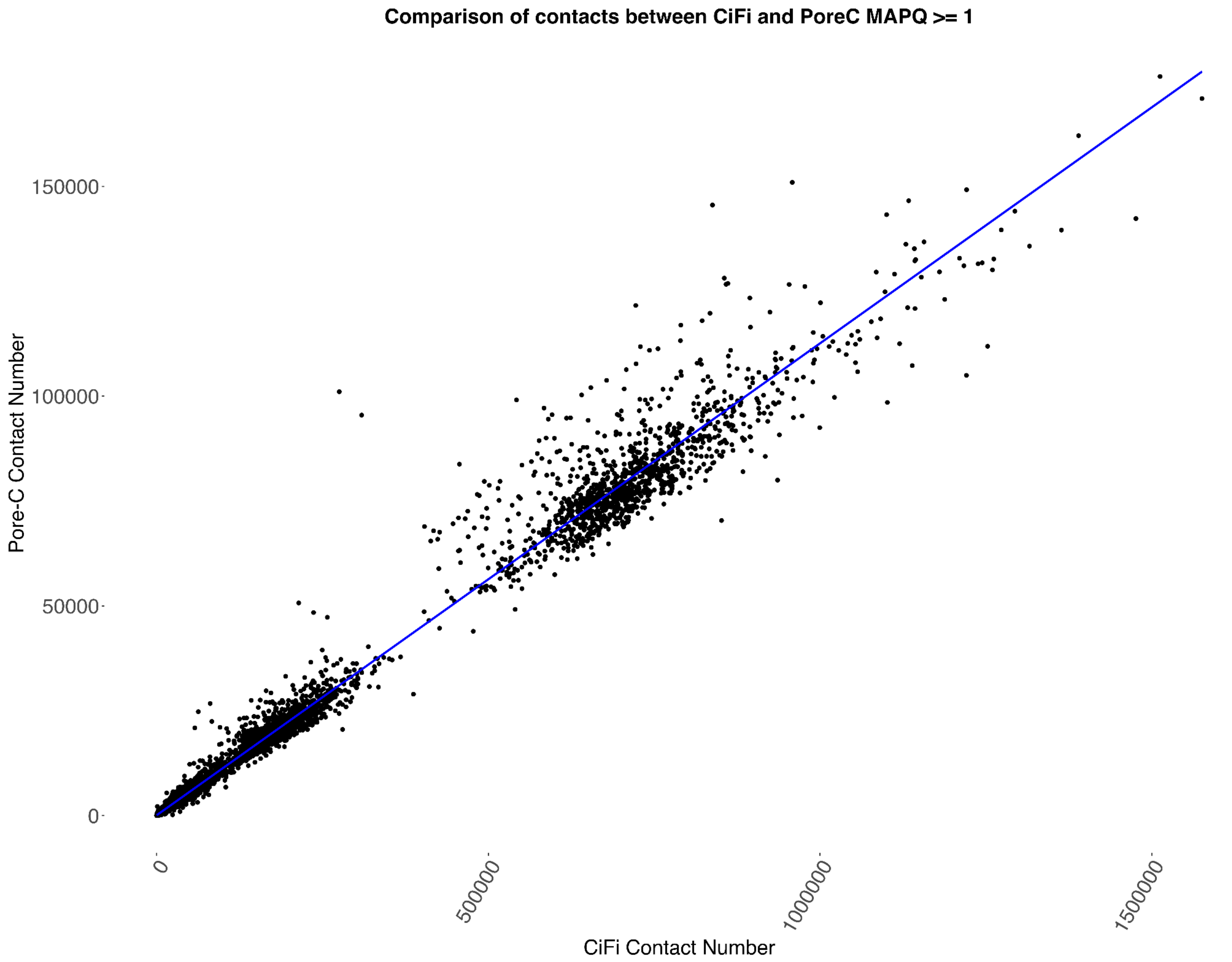

**Supplementary Figure 10. Correlation of chromatin contact signal at 2.5 Mbp resolution between DpnII CiFi and Pore-C.** $R^2 = 0.987$. These represent contact in 2.5-Mbp bins across the genome. Source data are provided as a Source Data file.

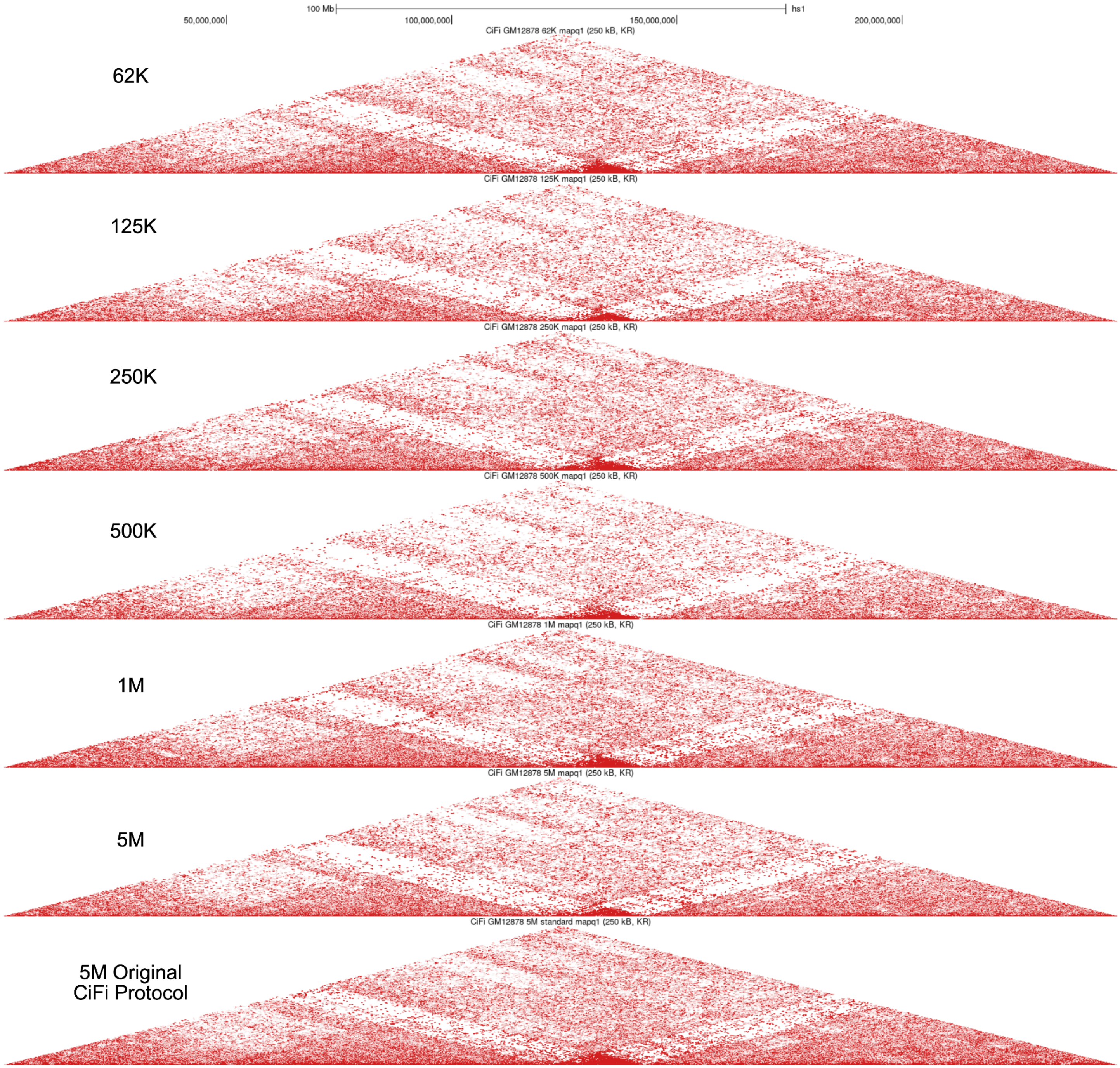

**Supplementary Figure 11. Comparisons of chromatin contacts for human LCL GM12878 across different cell titrations for chromosome 1.** Contacts are normalized using Knight–Ruiz algorithm and visualized at 250-kbp resolution.The starting cell amounts are listed on the left ranging from 62K to 5M. The 5M original CiFi protocol matches what was done for the initially successful CiFi run. Source data are provided through NCBI BioProject accession PRJEB83708.

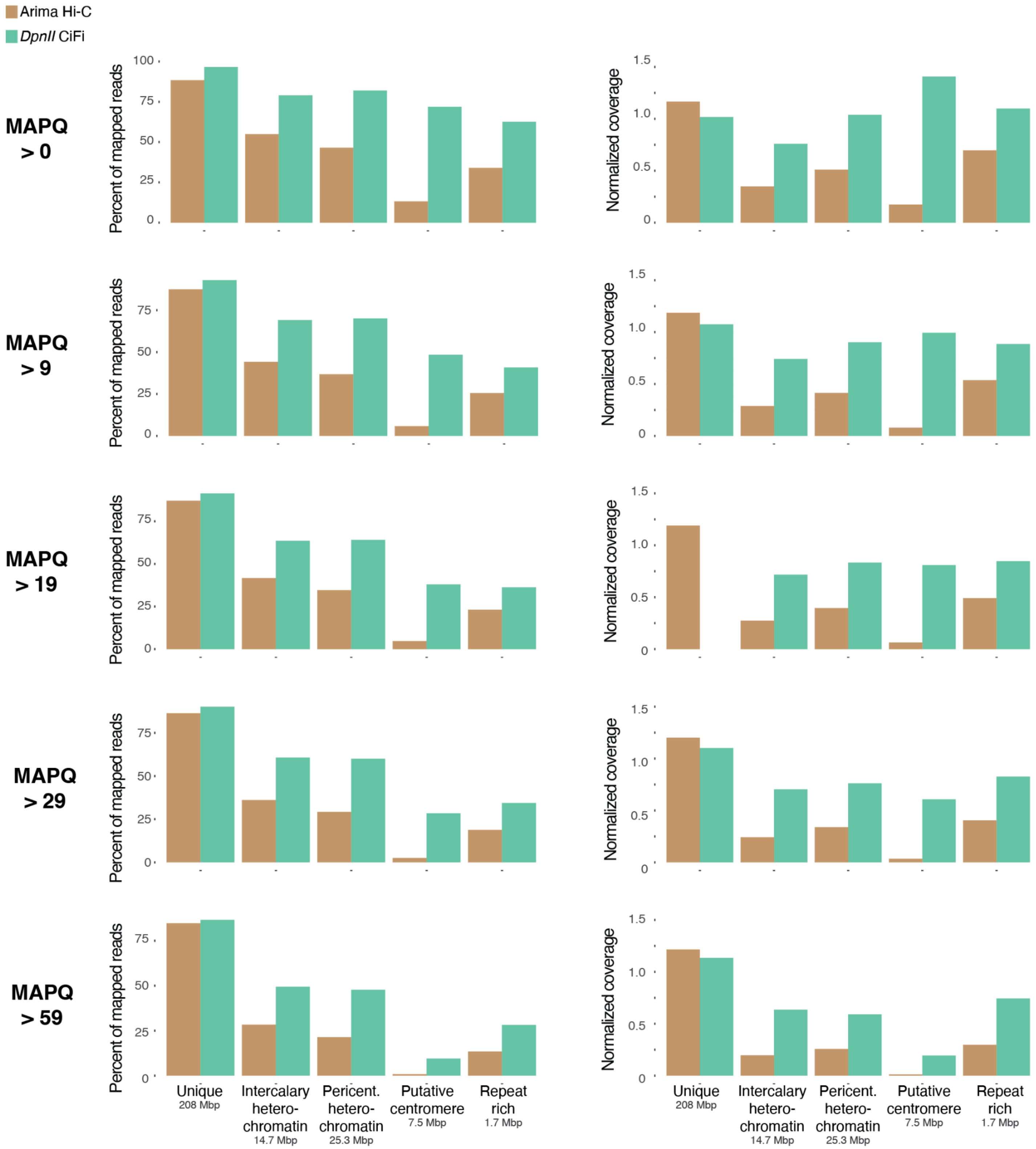

**Supplementary Figure 12. Percentage of reads mapped and normalized read coverage of CiFi and Hi-C across the genome for *Anopheles coluzzii*.** Shown on the left is percent of mapped reads at varied MAPQ cutoffs (indicated to the left of plots) of total mapped reads (with no MAPQ cutoff) per region. for Hi-C with Illumina (Arima v2) and CiFi DpnII across different repetitive genome classifications (intercalary heterochromatin, pericentromeric (pericent.) heterochromatin, putative centromeres, and repeat rich regions; see Supplementary Table 6) with total lengths of each classification type across the genome indicated. Shown on the right is read-coverage ratio (ratio of coverage per region to genomewide coverage) with varied MAPQ cutoffs for the same regions and sequencing libraries. Source data are provided as a Source Data file.

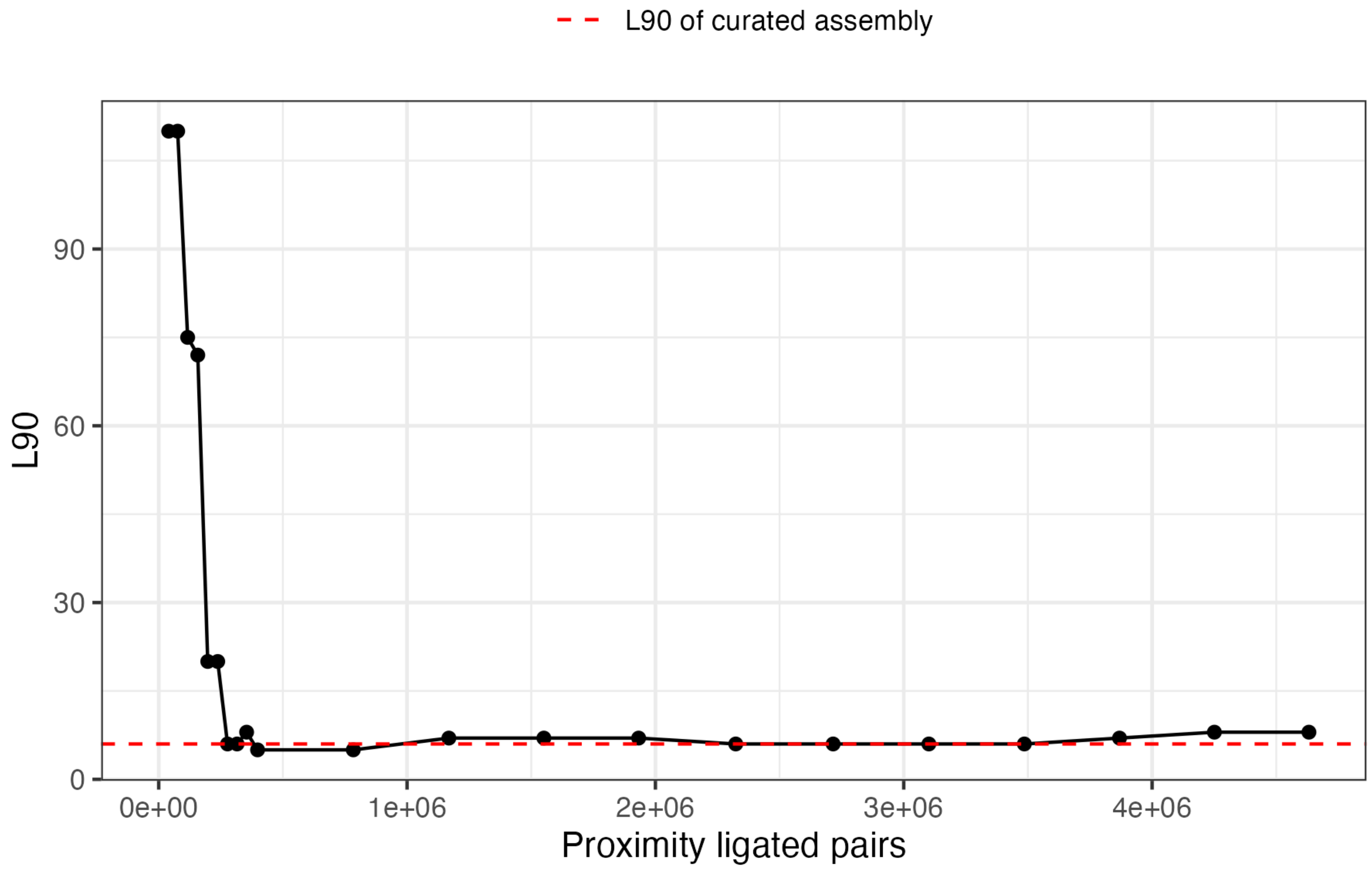

**Supplementary Figure 13. Scaffolding the Mediterranean fruit fly assembly with CiFi.** The number of proximity ligated pairs from the HindIII CiFi library necessary to scaffold a *C. capitata* genome. For this 600-Mbp phased genome, ~300,000 proximity-ligated pairs from 70,000 CiFi reads (~1.5x coverage of the genome) was necessary for scaffolding the phased assembly. A dashed red line denoting the L90 of haplotype one of a chromosome-scale reference assembly demonstrating the theoretical maximum L90 achievable at a chromosome-scale assembly. Source data are provided as a Source Data file.

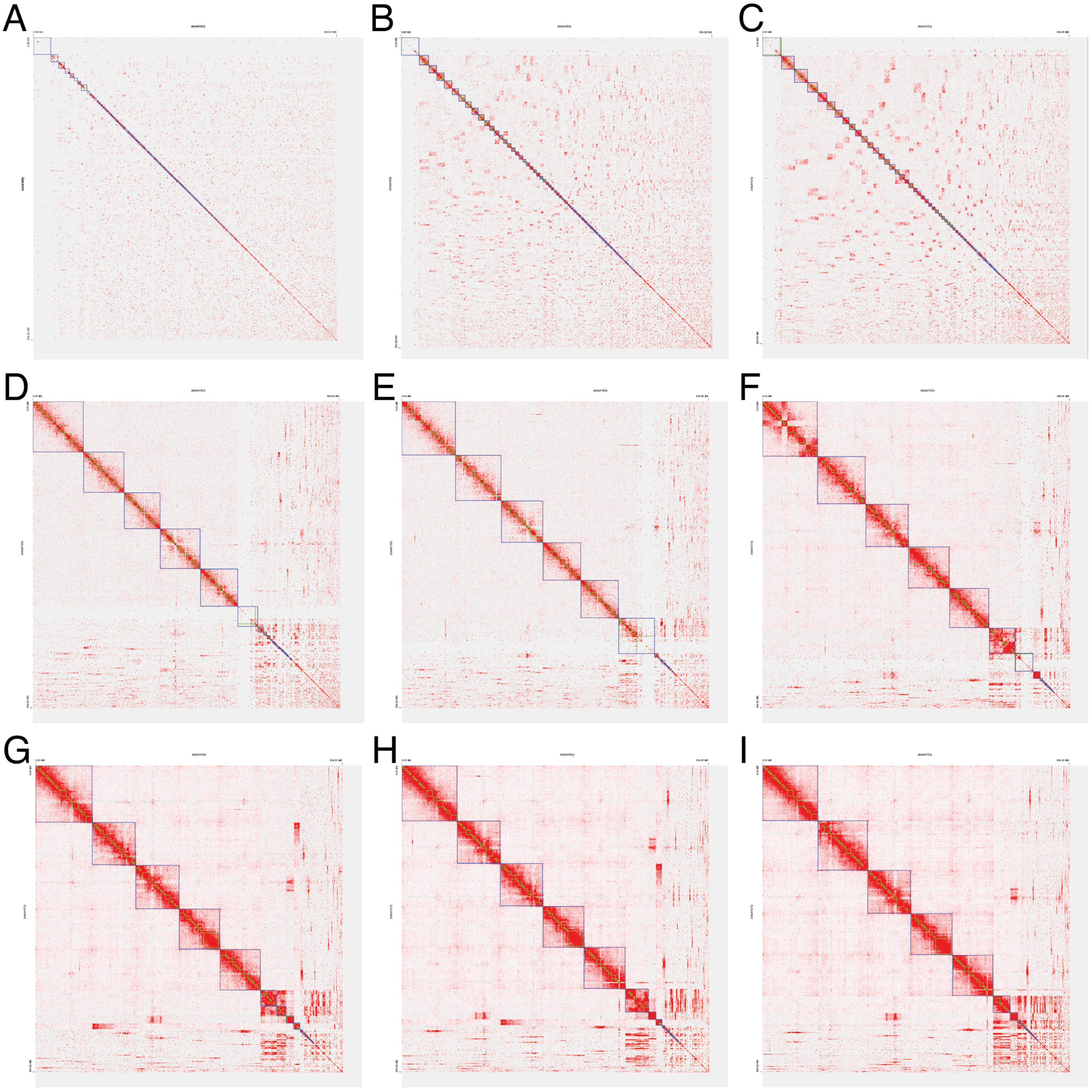

**Supplementary Figure 14. Scaffolding the Mediterranean fruit fly assembly with varying coverages of CiFi segments.** Subsampling paired segments of HindIII CiFi at (A) 10k read pairs, (B) 30k read pairs, (C) 50k read pairs, (D) 70k read pairs, (E) 100k read pairs, (F) 300k read pairs, (G) 500k read pairs, (H) 700k read pairs, and (I) 1000k read pairs produced varying number of chromosome scaffolds from YaHS. Source data are provided through NCBI BioProject accession PRJEB83708.

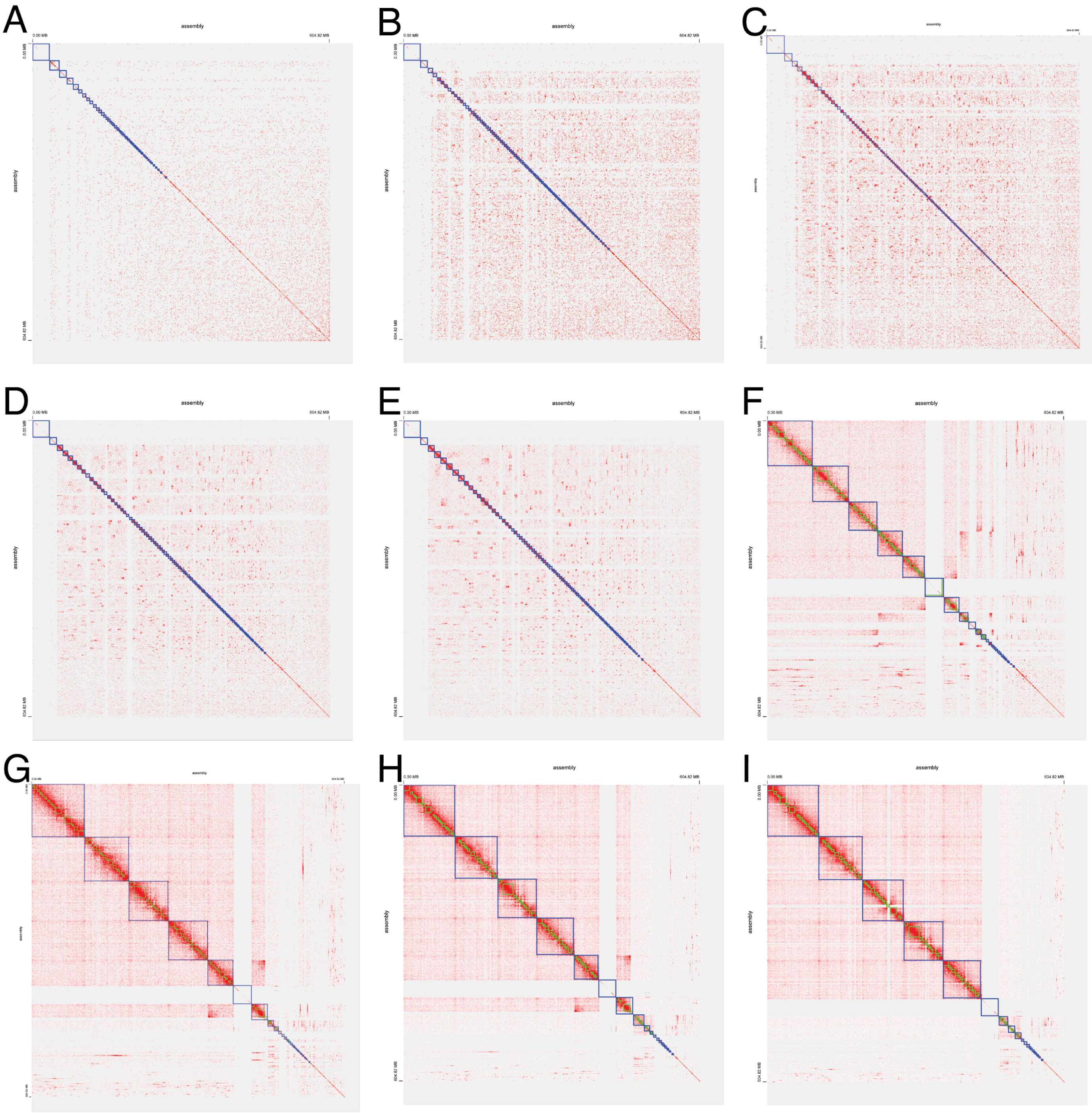

**Supplementary Figure 15. Scaffolding the Mediterranean fruit fly assembly with varying coverages of Hi-C reads.** Subsampling paired short reads (Element Biosciences Aviti) of Hi-C with DdeI and DpnII at (A) 10k read pairs, (B) 30k read pairs, (C) 50k read pairs, (D) 70k read pairs, (E) 100k read pairs, (F) 300k read pairs, (G) 500k read pairs, (H) 700k read pairs, and (I) 1000k read pairs produced varying number of chromosome scaffolds from YaHS. Source data are provided through NCBI BioProject accession PRJEB83708.

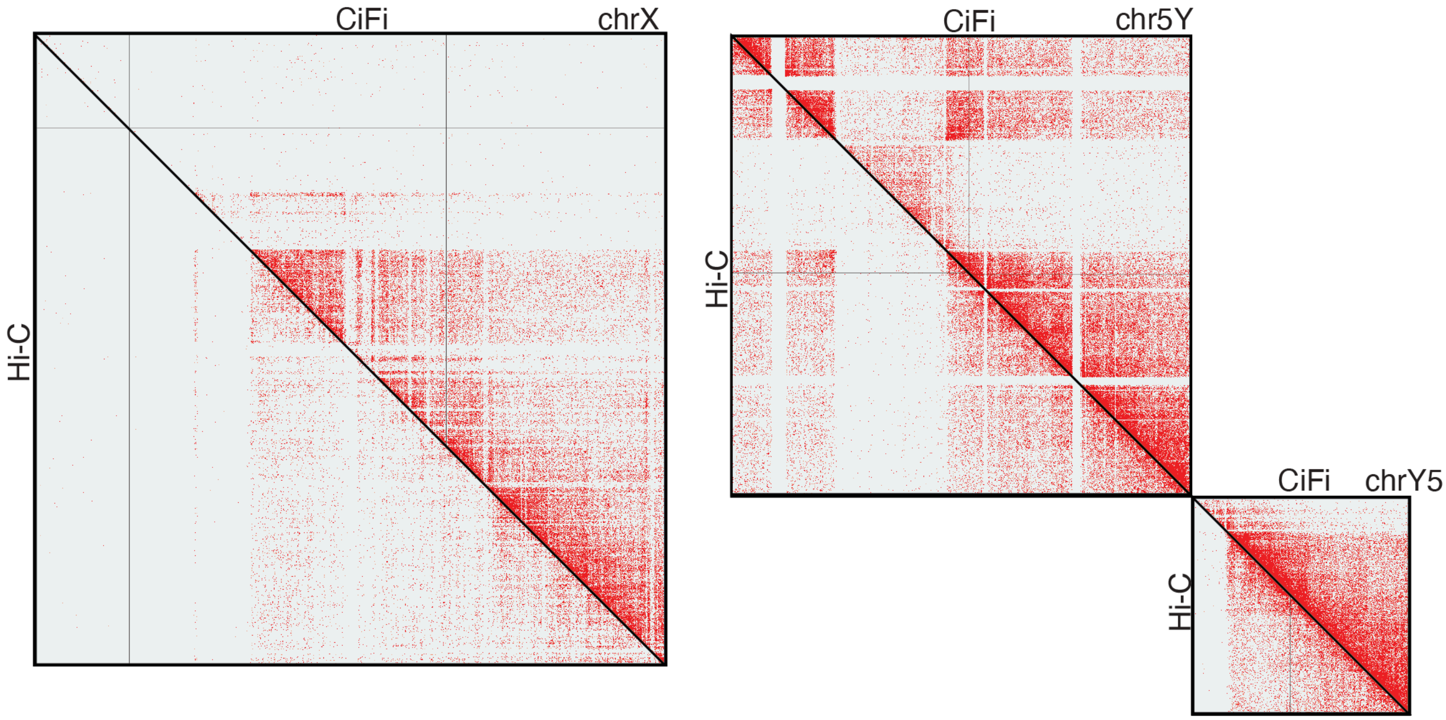

**Supplementary Figure 16. Comparison of CiFi and Hi-C datasets across sex chromosomes mapped to the new Mediterranean fruit fly genome assembly.** Pairwise contact comparisons between CiFi (top of diagonal) and Hi-C (bottom of diagonal) are visualized using Juicebox using a MAPQ cutoff > 0, 1-Mbp resolution, and Balanced ++ normalization, with notable dropouts in interactions in the Hi-C dataset. Source data are provided through NCBI BioProject accession PRJEB83708.

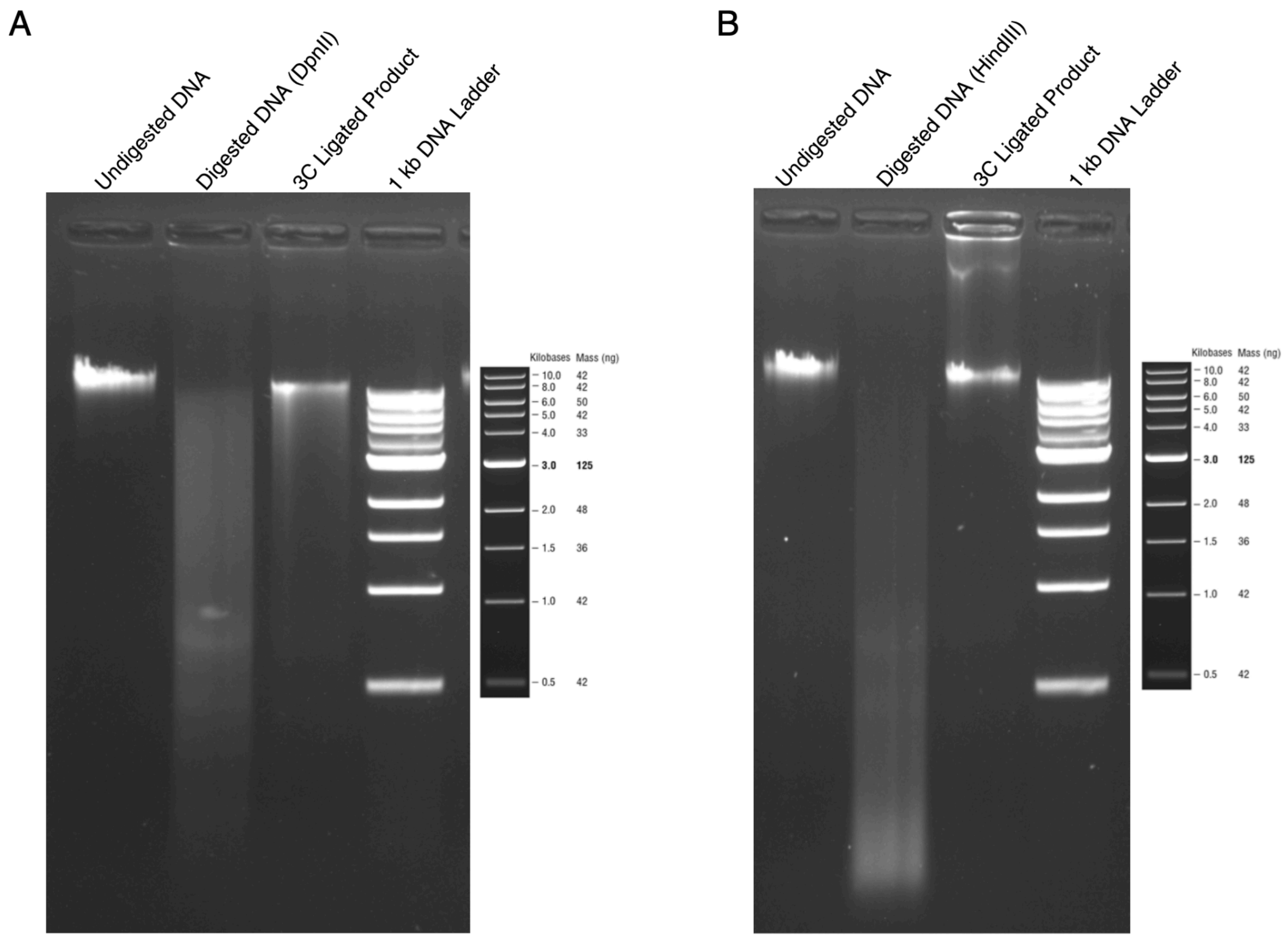

**Supplementary Figure 17. Quality check of 3C DNA.** 3C DNA obtained from GM12878 cells was analyzed on a 1% agarose gel alongside undigested and digested DNA samples for **(A)** DpnII and **(B)** HindIII accompanied with a 1 kbp DNA ladder as a molecular weight marker.

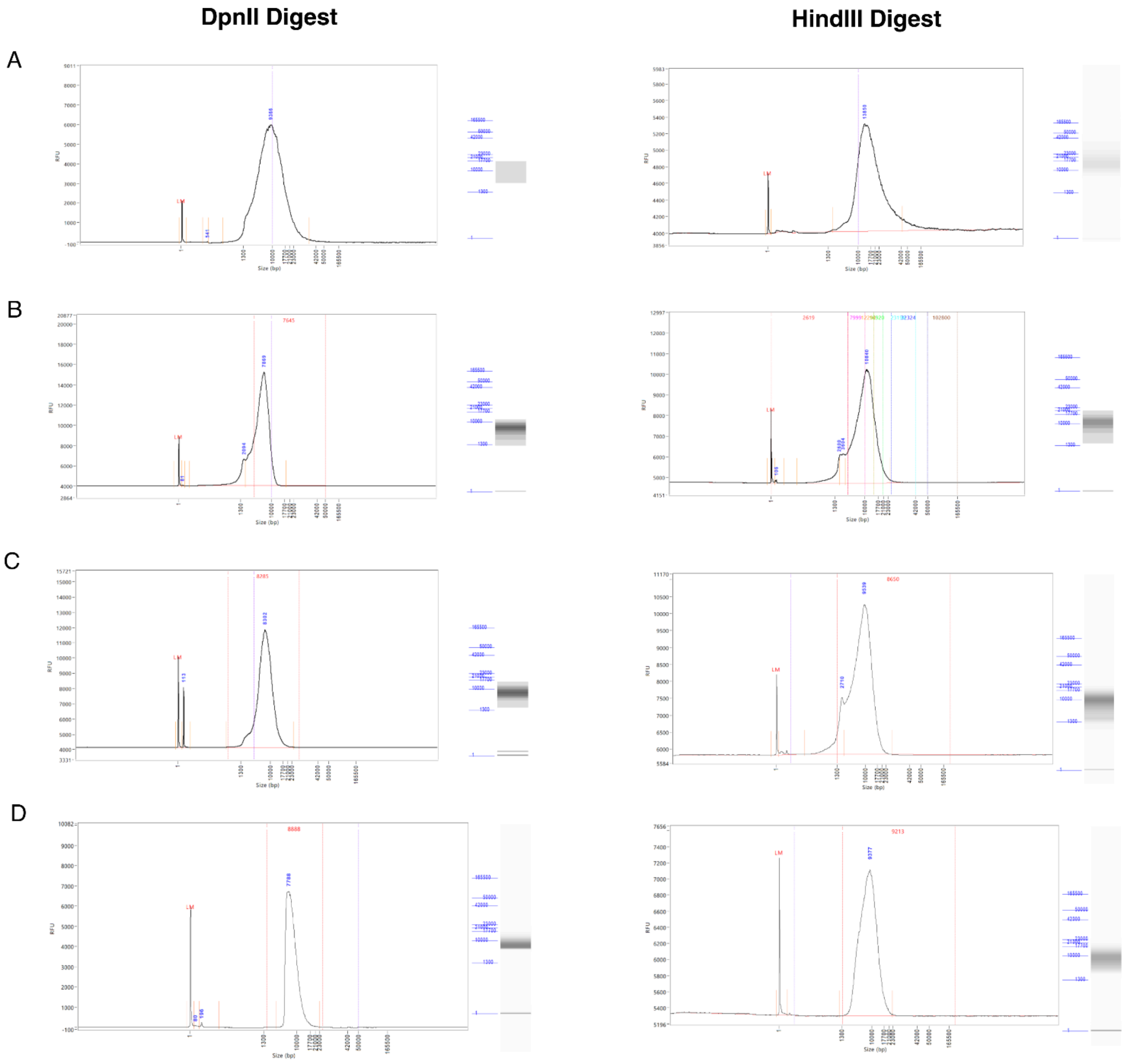

**Supplementary Figure 18. Verification of fragment size during CiFi library preparation.** Fragment size distributions were analyzed at key stages of the CiFi library preparation protocol using the Femto Pulse system (Agilent Technologies) for both DpnII (left column) and HindIII (right column) digests. The rows correspond to the following steps in the protocol: **(A)** 3C DNA QC, (**B)** After amplification, **(C)** After SMRTbell library preparation, and (**D)** After BluePippin size selection.

### 2. Supplementary Tables

**Supplementary Table 1. Comparison of Pore-C and Multi-Contact 3C protocols**

| Step | Pore-C Protocol | MC-3C Protocol |
|---|---|---|
| **Reference** | https://www.nature.com/articles/s41587-022-01289-z [2] | https://www.nature.com/articles/s41594-020-0506-5 [3] |
| **Cell Washing** | 10 million cells, washed three times in chilled 1X PBS, pelleted at 500xg for 5 min at 4°C. | 5 million cells, washed with Hank's balanced salt solution, pelleted. |
| **Cross-linking** | 1% formaldehyde, 10 min at room temperature, quenched with glycine to 125 mM. | 1% formaldehyde, 10 min at room temperature, quenched with glycine to 125 mM. |
| **Post-Crosslinking** | Incubated 5 min at room temperature, then 10 min on ice; pelleted at 500xg for 5 min at 4°C. | Incubated for 5 min at room temperature, followed by 15 min on ice. |
| **Cell Lysis** | Protease inhibitor in permeabilization buffer (10 mM Tris-HCl, 10 mM NaCl, 0.2% IGEPAL), 15 min on ice. | Dounce homogenizer in lysis buffer (10 mM Tris-HCl, 10 mM NaCl, 0.2% IGEPAL) for 15 min on ice. |
| **SDS Denaturation** | 1% SDS, incubated at 65°C for 10 min, quenched with 1% Triton X-100, 10 min on ice. | 0.1% SDS, incubated at 65°C for 10 min, quenched with Triton X-100 to 1%. |
| **Digestion** | DpnII (1 U/µL), incubated at 37°C for 18 hours with gentle rotation (<1000 rpm). | DpnII (400 U), incubated overnight at 37°C. |
| **Enzyme Inactivation** | Heat inactivation at 65°C for 20 min with rotation. | Heat inactivation at 65°C for 20 min. |
| **Proximity Ligation** | T4 ligase (100 µL buffer, 50 µL enzyme), incubated at 16°C for 6 hours with gentle rotation. | T4 ligase, incubated at 16°C for 4 hours. |
| **Crosslink Reversal** | Proteinase K (20 mg/mL) and SDS, incubated at 56°C for 18 hours with gentle rotation. | Proteinase K, incubated at 65°C for 2 hours, followed by overnight incubation at 65°C. |
| **DNA Purification** | Phenol-chloroform extraction, ethanol precipitation, SPRI bead size selection (>1.5 Kbp). | Phenol-chloroform extraction, ethanol precipitation, RNase treatment, size selection with BluePippin. |
| **Sequencing Platform** | Oxford Nanopore (MinION, GridION, PromethION). | PacBio RS II |

**Supplementary Table 2. Contact signal correlation for GM12878**

| Resolution | CiFi vs Illumina | CiFi vs Pore-C |
|---|---|---|
| 2500000 | 88.69 | 98.65 |
| 1000000 | 84.49 | 94.39 |
| 500000 | 82.55 | 96.25 |
| 250000 | 84.72 | 92.25 |
| 100000 | 80.52 | 87.45 |
| 50000 | 78.74 | 84.77 |
| 25000 | 73.89 | 73.65 |
| 10000 | 58.15 | 50.37 |

Correlations represented as $R^2$ values * 100

**Supplementary Table 3. Genome-wide TopDom TAD measure of concordance**

| | CiFi vs Illumina | CiFi vs Porce-C |
|---|---|---|
| **Chrom** | **MoC All** | **MoC All** |
| chr1 | 0.818 | 0.787 |
| chr2 | 0.831 | 0.795 |
| chr3 | 0.853 | 0.789 |
| chr4 | 0.824 | 0.763 |
| chr5 | 0.828 | 0.817 |
| chr6 | 0.834 | 0.784 |
| chr7 | 0.850 | 0.805 |
| chr8 | 0.837 | 0.820 |
| chr9 | 0.720* | 0.767* |
| chr10 | 0.836 | 0.800 |
| chr11 | 0.814 | 0.793 |
| chr12 | 0.846 | 0.805 |
| chr13 | 0.822 | 0.795 |
| chr14 | 0.820 | 0.772 |
| chr15 | 0.849 | 0.777 |
| chr16 | 0.800 | 0.785 |
| chr17 | 0.822 | 0.834 |
| chr18 | 0.819 | 0.772 |
| chr19 | 0.790 | 0.792 |
| chr20 | 0.844 | 0.814 |
| chr21 | 0.777 | 0.734 |
| chr22 | 0.758 | 0.758 |
| chrX | 0.774 | 0.745 |
| Genome-wide | 0.820 | 0.789 |

* Chr9 concordance generated from non-normalized contact matrices because a 27 Mbp Hsat causes Knight–Ruiz normalization to fail

**Supplementary Table 4. Genome-wide TopDom TAD Jaccard Index**

| | CiFi vs Illumina | | CiFi vs Pore-C | |
|---|---|---|---|---|
| Chrom | Jaccard Index | Jaccard Index 25% reciprocal | Jaccard Index | Jaccard Index 25% reciprocal |
| chr1 | 0.866 | 0.568 | 0.936 | 0.681 |
| chr2 | 0.973 | 0.744 | 0.984 | 0.768 |
| chr3 | 0.981 | 0.750 | 0.968 | 0.824 |
| chr4 | 0.974 | 0.606 | 0.968 | 0.637 |
| chr5 | 0.966 | 0.705 | 0.986 | 0.519 |
| chr6 | 0.985 | 0.609 | 0.986 | 0.556 |
| chr7 | 0.977 | 0.705 | 0.973 | 0.604 |
| chr8 | 0.968 | 0.783 | 0.989 | 0.415 |
| chr9 | 0.952* | 0.581* | 0.940* | 0.504* |
| chr10 | 0.974 | 0.930 | 0.970 | 0.752 |
| chr11 | 0.953 | 0.772 | 0.973 | 0.566 |
| chr12 | 0.973 | 0.871 | 0.965 | 0.571 |
| chr13 | 0.978 | 0.677 | 0.963 | 0.254 |
| chr14 | 0.957 | 0.708 | 0.968 | 0.818 |
| chr15 | 0.962 | 0.805 | 0.884 | 0.756 |
| chr16 | 0.847 | 0.667 | 0.939 | 0.727 |
| chr17 | 0.957 | 0.730 | 0.979 | 0.778 |
| chr18 | 0.941 | 0.681 | 0.974 | 0.825 |
| chr19 | 0.904 | 0.538 | 0.956 | 0.811 |
| chr20 | 0.945 | 0.898 | 0.964 | 0.546 |
| chr21 | 0.958 | 0.858 | 0.956 | 0.676 |
| chr22 | 0.876 | 0.575 | 0.941 | 0.803 |
| chrX | 0.966 | 0.588 | 0.987 | 0.733 |
| Genome-wide | 0.953 | 0.702 | 0.965 | 0.649 |

* Chr9 concordance generated from non-normalized contact matrices because a 27-Mbp Hsat causes KR normalization to fail

**Supplementary Table 5. Correlation of CiFi chromatin contacts across cell titration**

| Cell amount | Pairwise matrix resolution |
|---|---|
| 62K | 96.89 |
| 125K | 97.94 |
| 250K | 99.28 |
| 500K | 98.62 |
| 1M | 99.39 |
| 5M | 99.32 |

Correlations at 2.5 Mbp resolution represented as $R^2$ values * 100 versus 10M cell preparation

**Supplementary Table 6. Genomic coordinates of AcolN3 repeat annotations**

| Region | Chrom | Start | Stop |
|---|---|---|---|
| Intercalary Heterochromatin | chr2 | 72660000 | 81980000 |
| | chr3 | 38460000 | 40730000 |
| | chr3 | 56220000 | 59360000 |
| Pericentric Heterochromatin | chr2 | 57030000 | 70120000 |
| | chr3 | 50500000 | 54300000 |
| | chrX | 19890000 | 28269272 |
| Putative Centromere | chr2 | 61280556 | 67127712 |
| | chr3 | 53978800 | 54025408 |
| | chrX | 26627196 | 28269272 |
| Repeat Rich | chr2 | 84560000 | 85140000 |
| | chr3 | 23290000 | 23580000 |
| | chr3 | 61360000 | 62160000 |

**Supplementary Table 7.** ***An. coluzzii*** **mosquito chromosome coverage at MAPQ 1**

| | CiFi | | | | | Hi-C | | | | |
|---|---|---|---|---|---|---|---|---|---|---|
| Region | length | bases | mean | min | max | length | bases | mean | min | max |
| chr X | 28269272 | 789188012 | 27.92 | 0 | 27148 | 28269272 | 1696383258 | 60.01 | 0 | 915 |
| chr 2 | 126027569 | 6768633425 | 53.71 | 0 | 21762 | 126027569 | 8095043006 | 64.23 | 0 | 755 |
| chr 3 | 95248607 | 5372361774 | 56.40 | 0 | 24603 | 95248607 | 6347882665 | 66.65 | 0 | 3475 |
| Unique | 207876176 | 10906625462 | 52.47 | 0 | 2700 | 207876176 | 17987877261 | 86.53 | 0 | 1201 |
| Intercalary heterochromatin | 14730000 | 576689242 | 39.15 | 0 | 21762 | 14730000 | 381465781 | 25.90 | 0 | 1164 |
| Pericentromeric heterochromatin | 25269272 | 1352218895 | 53.51 | 0 | 27148 | 25269272 | 956938062 | 37.87 | 0 | 3570 |
| Putative centromere | 7535840 | 546707579 | 72.55 | 0 | 1019 | 7535840 | 97406319 | 12.93 | 0 | 3570 |
| Repeat rich | 1670000 | 94649612 | 56.68 | 0 | 6877 | 1670000 | 86257448 | 51.65 | 0 | 11078 |

**Supplementary Table 8. Contact signal correlation for *An. coluzzii* mosquito between CiFi and Hi-C**

| Resolution | CiFi vs Hi-C $R^2$ |
|---|---|
| 2500000 | 89.31 |
| 1000000 | 87.02 |
| 500000 | 91.09 |
| 250000 | 91.05 |
| 100000 | 88.77 |
| 50000 | 82.00 |
| 25000 | 71.40 |
| 10000 | 51.18 |

Correlations represented as $R^2$ values * 100

**Supplementary Table 9. Mediterranean fruit fly assembly statistics**

| | HindIII hap1 | HindIII hap2 | NlaIII hap1 | NlaIII hap2 |
|---|---|---|---|---|
| **Total length (Mb)** | 608.172 | 452.658 | 580.245 | 470.770 |
| **Contigs** | 443 | 453 | 453 | 320 |
| **Scaffolds** | 164 | 86 | 159 | 74 |
| **Contig N50 (Mb)** | 4.637 | 2.291 | 6.188 | 4.615 |
| **Contig L50** | 35 | 45 | 30 | 30 |
| **Scaffold N50 (Mb)** | 98.717 | 81.366 | 83.758 | 84.829 |
| **Scaffold L50** | 3 | 3 | 3 | 3 |
| **Contig N90 (Mb)** | 0.822 | 0.433 | 0.995 | 0.891 |
| **Contig L90** | 146 | 201 | 119 | 110 |
| **Scaffold N90 (Mb)** | 72.643 | 77.855 | 68.133 | 81.571 |
| **Scaffold L90** | 6 | 5 | 6 | 5 |
| **Longest contig (Mb)** | 35.237 | 14.57 | 16.094 | 17.772 |
| **Longest scaffold (Mb)** | 112.907 | 112.402 | 115.539 | 113.583 |
| **Complete Diptera BUSCOs (out of n = 3285)** | 99.3% | 96.7% | 99.4% | 98.1% |
| **Complete single-copy** | 97.0% | 95.8% | 98.3% | 97.7% |
| **Complete duplicated** | 2.3% | 0.9% | 1.1% | 0.4% |
| **Fragmented** | 0.2% | 0.3% | 0.3% | 0.2% |
| **Missing** | 0.5% | 3.0% | 0.3% | 1.7% |
| **Raw Quality Value** | 57.039 | 57.569 | 53.364 | 59.038 |
| **Adjusted Quality Value** | 58.098 | 55.777 | 41.374 | 58.32 |

### 3. Extended Experimental Procedures

#### Detailed CiFi Protocol

In this study, we report the development and results of the CiFi workflow, designed to capture high-resolution chromatin interactions using HiFi SMRTbell libraries prepared from ultra-low DNA input. The optimized protocol integrates chromatin conformation capture (Hi-C) with PacBio's SMRTbell sequencing, enabling efficient library preparation for long-read sequencing from limited DNA quantities. A detailed step-by-step protocol is provided below.

#### Part 1: 3C library preparation

**Note:** Please refer to Supplementary Data 3 for details on lower cell inputs.

We recommend keeping the samples on ice at all times when not incubating, unless stated otherwise in the protocol, and using wide-bore tips throughout the procedure.

##### i. Cross-linking and quenching

1. **Preparation**:
    - Wash 5-10 million cells three times in chilled 1X phosphate buffered saline (PBS) in a 50 mL centrifuge tube.
    - Centrifuge at 500xg for 5 minutes at 4°C between each wash.
    - Resuspend cells in 10 mL room temperature 1X PBS with 1% formaldehyde [EMD Millipore cat no. 818708] by gently pipetting **with a wide bore tip.**
    - Incubate at room temperature for 10 minutes.
2. **Quenching**:
    - Add 527 µL of 2.5 M glycine to achieve a final concentration of 1% (125 mM) in 10.5 mL.
    - Incubate for 5 minutes at room temperature, followed by 10 minutes on ice.
    - Pellet the cross-linked cells by centrifugation at 500xg for 5 minutes at 4°C.
    - Wash the cross-linked cells with 1x PBS, remove the supernatant, and snap-freeze the pellet in liquid nitrogen. Store at -80°C until you start the protocol.

##### ii. Restriction enzyme digestion

1. **Cell lysis**:
    - Resuspend the cell pellet in a mixture of 50 µL of protease inhibitor cocktail [Sigma Aldrich cat no. P8340] in 500 µL of cold permeabilization buffer (10 mM Tris-HCl pH 8.0, 10 mM NaCl, 0.2% IGEPAL CA-630, nuclease-free water).
    - Place on ice for 15 minutes.
    - Centrifuge at 500xg for 10 minutes at 4°C.
    - Aspirate the supernatant and replace with 200 µL of chilled 1.5X digestion reaction buffer [NEB] compatible with the restriction enzyme used.
    - Centrifuge again at 500xg for 10 minutes at 4°C, then aspirate and resuspend in 300 µL of chilled 1.5X digestion reaction buffer.

2. **Chromatin denaturation**:
    - Add 33.5 µL of 1% (w/v) SDS [Thermo Fisher Scientific cat no. 15553027] to each cell suspension.
    - Incubate for exactly 10 minutes at 65°C with gentle agitation.
    - Place on ice immediately afterwards.

- Quench the SDS by adding 37.5 µL of 10% (v/v) Triton X-100 [Sigma Aldrich cat no. 93443] for a final concentration of 1%.
- Incubate for 10 minutes on ice.

3. **Digestion**:
   - Permeabilized cells are then digested with a final concentration of 1 U/µL of DpnII [NEB], brought to volume with nuclease-free water to achieve a final 1X digestion reaction buffer in 450 µL.
   - Mix by gentle inversion and incubate in a thermomixer at 37°C for 18 hours with periodic < 1000 rpm rotation (< 30 seconds every 15 minutes) to prevent condensation inside the lid.

**iii. Proximity ligation and reverse cross-linking**

1. **Inactivation**:
   - DpnII restriction digests are heat-inactivated at 65°C for 20 minutes with 300 rpm rotation. Place on ice immediately afterwards.
2. **Ligation**:
   - Set up proximity ligation at room temperature with the following reagents:
     - 100 µL of 10X T4 DNA ligase buffer [NEB]
     - 10 µL of 10 mg/mL BSA
     - 50 µL of T4 DNA ligase [NEB M0202L]
     - **Total volume of 1000 µL** with nuclease-free water.
   - Cool to 16°C and incubate for 6 hours with gentle rotation.

**iv. Protein degradation and DNA purification**

1. **Reverse cross-linking**:
   - Treat samples with:
     - 100 µL 20 mg/mL Proteinase K [Thermo Fisher Scientific cat no. 25530049]
     - 100 µL 10% SDS [Thermo Fisher Scientific cat no. 15553027]
     - 500 µL 20% (v/v) Tween-20 [Sigma Aldrich cat no. P9416]
     - **Total volume of 2000 µL** with nuclease-free water.
   - Incubate in a thermomixer at 56°C for 18 hours with < 1000 rpm rotation (< 30 seconds every 15 minutes) to prevent condensation inside the lid.

2. **Purification**:
   - Transfer the sample to a 15 mL centrifuge tube, rinsing the original tube with a further 200 µL of nuclease-free water to collect any residual sample, bringing the total sample volume to 2.2 mL.
   - Purify DNA using standard phenol-chloroform extraction and ethanol precipitation methods.
   - Before proceeding, check undigested, digested, and ligated DNA products on an agarose gel to ensure the experiment was successful (Supplementary Figure 17).
   - Store the purified libraries at 4°C for short-term storage, and at -20°C for long-term storage until the next step.

**Size selection (AMPure PB beads)**

Follow the PacBio protocol for size selection using AMPure PB beads (PacBio, Cat# 100-265-900) at a 0.45× ratio for an ~3 kb cutoff, or using 35% diluted AMPure PB beads at a 3.1× ratio for an ~4-5 kb cutoff. Add 50 µL of TE buffer to resuspend the beads, then proceed to the next step.

- For libraries prepared using **DpnII**, proceed directly to size selection.
- For libraries prepared using **HindIII**, the longer DNA fragments require an additional shearing step before size selection. Shear the 3C DNA to approximately 10 kb using a g-TUBE (Covaris, 520104), following the manufacturer's protocol.
- After size selection, use 0.5 ng/µL of DNA for quality control (QC) with Femto Pulse automated pulsed-field capillary electrophoresis to confirm that the DNA matches the expected size distribution (Supplementary Figure 18A)

**Part 2: SMRTbell library preparation from modified ultra-low DNA input**

**Required PacBio kits:**
SMRTbell gDNA amplification kit, cat# 101-980-000
SMRTbell adapter index plate 96A, cat# 102-009-200
SMRTbell cleanup beads, cat# 102-158-300
SMRTbell express template prep kit 2.0, cat# 100-938-900*
Alternative Kit: SMRTbell prep kit 3.0, cat# 102-182-700*

*Note: The initial optimization of the CiFi protocol for GM12878 with DpnII used the SMRTbell Express Template Prep Kit 2.0, with the detailed protocol described below in Part 2A. CiFi libraries for GM12878 with HindIII, mosquito, and Mediterranean fruit fly used the SMRTbell Prep Kit 3.0, with the detailed protocol provided in Part 2B.

**For PCR amplification:**
KOD Xtreme hot-start polymerase from Sigma, cat# 71975-M

PacBio library construction requirements for ultra-low DNA input samples:

| Required gDNA Input Amount | Required Quality of Input gDNA | gDNA Shearing Method | Target Sheared Fragment Size Distribution Mode | Amplification Target Size Distribution Mode | Total Mass of Pooled PCR Product Required for Library Construction | Required SMRTbell Library Input for BluePippin Size-Selection |
|---|---|---|---|---|---|---|
| 5–20 ng | Majority of gDNA >20 kb | g-TUBE | 10 kb sheared DNA is optimal | 8–10 kb | ≥500 ng | ≥400 ng |

**Note:** Skip the gDNA shearing step for DpnII since we already digested it with the restriction enzyme, perform only for HindIII libraries (as described above).

**Part 2A: SMRTbell library preparation with Express Template Prep Kit 2.0**

**i. Removing single-strand overhangs**

1. **Preparation of DNA prep additive:**
    - Dilute the DNA prep additive with enzyme dilution buffer. Mix well and quickly spin.
        - 4.0 µL Enzyme dilution buffer
        - 1.0 µL DNA prep additive (stock)
        - **Total volume: 5 µL**
2. **Reaction mix preparation:** For each sample, prepare the following reaction mix in a PCR tube:
    - 7.0 µL DNA prep buffer
    - <45 µL 3C DNA
    - 0.6 µL NAD
    - 1.0 µL Diluted DNA prep additive (prepared above)
    - 1.0 µL DNA prep enzyme
    - $H_2O$ up to a **total volume of 55.0 µL**
3. Mix thoroughly using wide-bore pipette tips (pipette mix 10 times).
4. Quick spin to collect contents in a microfuge.
5. Place in a thermocycler and run the following program:
    - 15 minutes at 37°C
    - Hold at 4°C
6. Proceed to the next step.

**ii. Repair DNA damage**

1. **Reaction setup:** Add 2 µL DNA damage repair mix v2 directly to the reaction:
    - 55.0 µL Reaction mix
    - 2.0 µL DNA damage repair mix v2
    - **Total volume: 57.0 µL**
2. Mix thoroughly using wide-bore pipette tips (pipette mix 10 times).
3. Quick spin to collect contents in a microfuge.
4. Place in a thermocycler and run the following program:
    - 30 minutes at 37°C
    - Hold at 4°C
5. Proceed to the next step.

**iii. Repair ends/A-tailing**

1. **Reaction setup:** Add 3 µL end prep mix directly to the reaction:
    - 57.0 µL Reaction mix (damage-repaired sample)
    - 3.0 µL End prep mix
    - **Total volume: 60.0 µL**
2. Mix thoroughly using wide-bore pipette tips (pipette mix 10 times).
3. Quick spin to collect contents in a microfuge.
4. Place in a thermocycler and run the following program:
    - 30 minutes at 20°C
    - 30 minutes at 65°C
    - Hold at 4°C

5. Proceed to the next step.

**iv. Adapter ligation**

1. **Preparation of amplification adapters:** Dilute the Amplification Adapters with Duplex Buffer:
   - 9.0 µL Duplex Buffer
   - 1.0 µL Amplification Adapters
   - **Total volume: 10.0 µL** Use immediately.
2. **Reaction setup:** Add the following components to the reaction:
   - 60.0 µL Reaction mix (A-tail sample)
   - 2.5 µL Diluted amplification adapters
   - 30.0 µL Ligation mix
   - 1.0 µL Ligation additive
   - 1.0 µL Ligation enhancer
   - **Total volume: 94.5 µL**
3. Mix thoroughly using wide-bore pipette tips (pipette mix 10 times).
4. Quick spin to collect contents in a microfuge.
5. Place in a thermocycler and run the following program:
   - 60 minutes at 20°C
   - Hold at 4°C
6. Proceed to the next step.

**v. Purification of SMRTbell library**

1. Bring SMRTbell beads (PacBio) to room temperature for 30 to 60 minutes before use.
2. Add 77.5 µL SMRTbell beads to 94.5 µL Reaction Mix. Mix gently (pipette mix 10 times).
3. Incubate on the bench for 5 minutes at room temperature.
4. Place the tube on a magnetic stand and wait until the supernatant is clear. Remove the supernatant.
5. Wash beads twice with 200 µL freshly prepared 80% ethanol.
6. After the second ethanol wash, spin briefly, return to the magnetic stand, and remove residual ethanol. Do not let the beads dry out.
7. Resuspend beads in 97 µL of EB, pipette mix 10 times, and incubate at 37°C for 10 minutes to elute DNA.
8. Place on a magnetic stand to separate beads. Transfer 97 µL of purified sample to a new tube and set aside on ice.

IMPORTANT: For the Library Amplification by PCR, the 97 µL of the purified eluted sample will be divided and used for three reactions to achieve enough output product for the next step. Each reaction requires a volume of 32 µL of the purified eluted sample. Sometimes, we need to repeat the PCR reaction 2x or 3x to achieve 1 µg of output.

**vi. Library amplification by modified PCR**

1. **Reaction setup:** For each reaction:
   - 60 µL 2X Xtreme buffer
   - 24 µL 2 mM dNTPs
   - 2.4 µL Sample amplification PCR primer
   - 32 µL DNA
   - 2.4 µL KOD Xtreme Hot Start DNA polymerase

     **Total: 120.8 µL**
2. **Thermal cycler settings:**
   - Initial Denaturation: 94°C for 2 minutes (1 cycle)

- Denaturation: 98°C for 10 seconds (13 cycles)
- Annealing: 60°C for 30 seconds (13 cycles)
- Extension: 68°C for 10 minutes (13 cycles)
- Final Extension: 68°C for 5 minutes (1 cycle)
- Hold: 4°C indefinitely

**vii. Purification of amplified DNA**

1. Bring SMRTbell beads to room temperature for 30 to 60 minutes before use.
2. Add 99 µL SMRTbell beads to the 120.8 µL reaction mix. Mix gently (pipette mix 10 times).
3. Incubate on the bench for 5 minutes at room temperature.
4. Place the tube on a magnetic stand and wait until the supernatant is clear. Remove the supernatant.
5. Wash beads twice with 200 µL freshly prepared 80% ethanol.
6. After the second ethanol wash, spin briefly, return to the magnetic stand, and remove residual ethanol. Do not let the beads dry out.
7. Resuspend beads in 26 µL of EB, pipette mix 10 times, and incubate at room temperature for 5 minutes to elute DNA.
8. Place on a magnetic stand to separate beads. Transfer 26 µL of eluted amplified DNA to a new tube and set aside on ice.
9. Use 1 µL of the sample to quantify with Qubit dsDNA HS kit.
10. Use 1 µL of amplified DNA for DNA sizing QC by Femto Pulse automated pulsed-field capillary electrophoresis (200-500 pg of sample) (Supplementary Figure 18B).
11. Store amplified DNA at 4°C or -20°C for future use.

**Total DNA requirement:**

- Ensure a pooled DNA mass ≥500 ng in 47.4 µL (recommended: 1 µg).

**viii. Repair DNA damage (Post-amplification)**

1. **Reaction Setup:** Add the following components to a single PCR tube:
    - 7.0 µL DNA Prep Buffer
    - ≤47.4 µL Pooled Amplified DNA
    - 0.6 µL NAD
    - 2.0 µL DNA Damage Repair Mix v2
    - $H_2O$ up to a **total volume of 57.0 µL**
2. Mix thoroughly using wide-bore pipette tips (pipette mix 10 times).
3. Quick spin to collect contents in a microfuge.
4. Place in a thermocycler and run the following program:
    - 30 minutes at 37°C
    - Hold at 4°C
5. Proceed to the next step.

**ix. Repair ends/A-tailing**

1. **Reaction setup:** Add 3 µL End Prep Mix directly to the reaction:
    - 57.0 µL Reaction mix (damage-repaired sample)
    - 3.0 µL End prep mix

- **Total volume: 60.0 µL**

2. Mix thoroughly using wide-bore pipette tips (pipette mix 10 times).
3. Quick spin to collect contents in a microfuge.
4. Place in a thermocycler and run the following program:
   - 30 minutes at 20°C
   - 30 minutes at 65°C
   - Hold at 4°C
5. Proceed to the next step.

**x. Adapter ligation**

1. **Reaction setup:** Add the following components to Reaction Mix (A-tail sample):
   - 60.0 µL Reaction mix
   - 5.0 µL Overhang adapter v3 (or SMRTbell adapter index plate for barcoding)
   - 30.0 µL Ligation mix
   - 1.0 µL Ligation additive
   - 1.0 µL Ligation enhancer
   - **Total volume: 97 µL**
2. Mix thoroughly using wide-bore pipette tips (pipette mix 10 times).
3. Quick spin to collect contents in a microfuge.
4. Place in a thermocycler and run the following program:
   - 60 minutes at 20°C
   - Hold at 4°C
5. Proceed to the next step: “Purification of SMRTbell Library.”

**xi. Purification of SMRTbell library**

1. Add 97 µL of room-temperature resuspended SMRTbell beads to the 97 µL reaction mix. Pipette mix 10 times.
2. Perform a quick spin to collect liquid.
3. Incubate on the bench for 5 minutes at room temperature.
4. Place on a magnetic stand to separate beads from supernatants. Remove supernatant using a P200 pipette.
5. Wash beads twice with 200 µL freshly prepared 80% ethanol. After the second wash, briefly spin and return to the magnet. Remove residual ethanol using a P20 pipette.
6. Resuspend beads in 32 µL EB, mix thoroughly, and incubate at room temperature for 5 minutes to elute DNA.
7. Place on the magnetic stand to separate beads. Transfer eluted DNA to a new tube.
8. Use 1 µL of SMRTbell library to quantify with Qubit dsDNA HS kit.
9. Use 1 µL for DNA sizing QC by Femto Pulse (200-500 pg sample) (Supplementary Figure 18C).
10. Proceed with "BluePippin Size-Selection of SMRTbell Library" or store at 4°C or -20°C.

**xii. BluePippin or diluted AMPure PB bead cleanup, size selection, and sequencing**

For size selection of SMRTbell libraries, you can use either BluePippin or diluted AMPure PB beads based on the desired fragment size and available resources. Both methods enrich DNA fragments within specific size ranges and are compatible with sequencing on the Revio or Sequel II platforms.

**- BluePippin size selection:**
Perform >5 kb size selection using BluePippin according to the manufacturer's protocol.

**- Alternatively, diluted AMPure PB bead cleanup and size selection:**
Perform ~3-5 kb size selection using diluted AMPure PB beads to enrich the sample for DNA fragments of the desired size.

**Instructions for diluted AMPure PB bead preparation and use:**

- **Prepare 35% AMPure PB beads:**
  Mix 1.75 mL of resuspended AMPure PB beads with 3.25 mL of elution buffer (35% v/v).
  Store the diluted beads at 4°C for up to 30 days.
- **Size selection procedure:**
  1. Add 3.1X volume of diluted beads to the sample.
  2. Mix thoroughly and briefly spin down.
  3. Incubate at room temperature for 10 minutes to bind DNA.
  4. Place on a magnetic rack and remove the supernatant.
  5. Wash beads twice with 200 µL of 80% ethanol.
  6. Air-dry the beads and resuspend in 50 µL of elution buffer.
  7. Incubate for 5 minutes and use the magnetic rack to separate the beads.
  8. Transfer the supernatant containing the size-selected DNA to a new tube for sequencing or storage.

After size selection, use 1 µL of the sample for library sizing QC by Femto Pulse (200-500 pg sample) (Supplementary Figure 18D)

**Part 2B: SMRTbell library preparation with Prep Kit 3.0**

**i. Repair and A-tailing of digested 3C DNA**

1. **Reaction setup: Add the following components to a PCR tube:**
   - 4 µL Repair buffer
   - 1 µL End repair mix
   - 0.5 µL DNA repair mix
   - 24.5 µL Digested 3C DNA (5-20 ng)
   - **Total volume: 30 µL**
2. Quick spin to collect contents in the tube.
3. Place in a thermocycler and run the following program:
   - 30 minutes at 37°C
   - 5 minutes at 65°C
   - Hold at 4°C
4. Proceed to the next step.

**ii. Ligation of linear amplification adapter and cleanup**

1. Reaction setup: Add the following components to a PCR tube:
   - 2 µL Amplification adapter
   - 10 µL Ligation mix
   - 0.5 µL Ligation enhancer
   - 30 µL End-prepared DNA
   - **Total volume: 42.5 µL**
2. Quick spin to collect contents in the tube.
3. Place in a thermocycler and run the following program:

    - 30 minutes at 20°C
    - Hold at 4°C
4. Perform cleanup using 1X SMRTbell cleanup beads as described in Part 2A. Add 24 µL of elution buffer to resuspend the beads, then proceed to the next step.

**iii. Amplification and cleanup**

1. Reaction setup: Add the following components to a PCR tube:
    - 50 µL 2X Xtreme buffer
    - 20 µL 2 mM dNTPs
    - 4 µL Sample amplification PCR primer
    - 2 µL KOD Xtreme Hot Start DNA polymerase
    - 24 µL Adapter-ligated sample
    - **Total volume: 100 µL**
2. Quick spin to collect contents in the tube.
3. Place in a thermocycler and run the following program:
    - Initial Denaturation: 94°C for 2 minutes (1 cycle)
    - Denaturation: 98°C for 10 seconds (10–12 cycles)
    - Annealing and Extension: 68°C for 10 minutes (10–12 cycles)
    - Final Extension: 68°C for 7 minutes (1 cycle)
    - Hold: 4°C indefinitely

**DNA input and PCR cycles:**

- 5 ng: 12 cycles
- 10 ng: 11 cycles
- 20 ng: 10 cycles

4. Perform cleanup using 1X SMRTbell cleanup beads as described in Part 2A. Add 50 µL of elution buffer to resuspend the beads. Quantify the DNA using the Qubit dsDNA HS Kit and check quality control (QC) using a Femto Pulse (Supplementary Figure 18B). Proceed to the next step.

**Total DNA requirement:**

- ≥500 ng of amplified DNA for Revio sequencing.
- ≥250 ng of amplified DNA for Sequel II/IIe sequencing

**iv. Repair and A-tailing of amplified DNA**

1. Reaction setup: Add the following components to a PCR tube:
    - 8 µL Repair buffer
    - 2 µL End repair mix
    - 1 µL DNA repair mix
    - 49 µL Amplified DNA
    - **Total volume: 60 µL**
2. Quick spin to collect contents in the tube.
3. Place in a thermocycler and run the following program:
    - 30 minutes at 37°C
    - 5 minutes at 65°C
    - Hold at 4°C

**v. SMRTbell adapter ligation and cleanup**

1. Reaction setup: Add the following components to a PCR tube:
   - 4 µL SMRTbell adapter (or SMRTbell adapter index plate for barcoding)
   - 15 µL Ligation mix
   - 1 µL Ligation enhancer
   - 60 µL Repaired DNA
   - **Total volume: 80 µL**
2. Quick spin to collect contents in the tube.
3. Place in a thermocycler and run the following program:
   - 30 minutes at 20°C
   - Hold at 4°C
4. Perform cleanup using 1X SMRTbell cleanup beads as described in Part 2A. Add 40 µL of elution buffer to resuspend the beads, then proceed to the next step.

**vi. Nuclease treatment**

1. Reaction Setup: Add the following components to a PCR tube:
   - 5 µL Nuclease buffer
   - 5 µL Nuclease mix
   - 40 µL Ligated DNA
   - **Total volume: 50 µL**
2. Place in a thermocycler and run the following program:
   - 15 minutes at 37°C
   - Hold at 4°C
3. Perform cleanup using 1X SMRTbell cleanup beads as described in Part 2A.

**vii. BluePippin or diluted AMPure PB bead cleanup, size selection, and sequencing**

As described in the previous section, Part 2A-xii, the procedure was followed to complete this step.

### 4. Uncropped Gel Image

**Supplementary Figure 17**

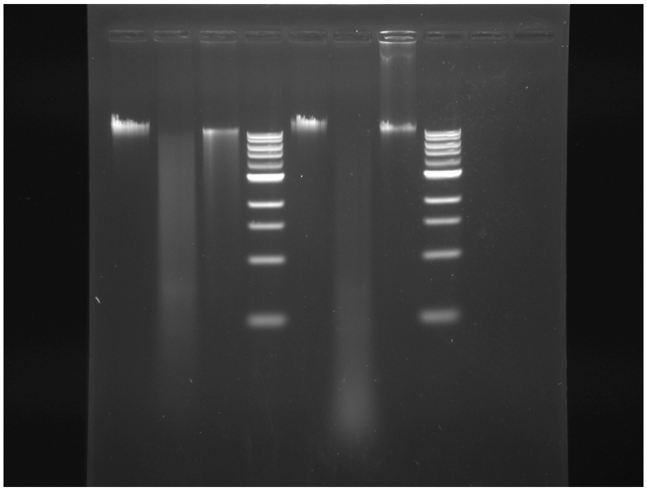
